# Selective Inactivation of Astrocytic Monoacylglycerol Lipase for Alzheimer’s Disease Therapy

**DOI:** 10.64898/2025.12.04.692105

**Authors:** Li Sun, Mei Hu, Jianlu Lyu, Dexiao Zhu, Fei Gao, Mingzhe Pan, Jian Zhang, Chu Chen

## Abstract

Alzheimer’s disease (AD) is the leading cause of dementia in the elderly, yet effective therapies remain lacking. Here, we identify monoacylglycerol lipase (MAGL), the principal enzyme that degrades the endocannabinoid 2-arachidonoylglycerol (2-AG) in the brain, specifically in astrocytes as a promising therapeutic target for AD. APP transgenic (TG) mice with astrocyte-specific MAGL deletion (TG-aKO), but not neuron-specific deletion (TG-nKO), exhibit markedly reduced Aβ pathology, neurodegeneration, and neuroinflammation, along with preserved synaptic structure and function and improved cognition. These benefits are recapitulated in TG mice by AAV-mediated silencing of astrocytic MAGL, whereas MAGL overexpression exacerbates neuropathology and accelerates synaptic and cognitive decline. Transcriptomic analyses reveal that dysregulated synapse-, inflammation-, and AD-related gene expression profiles are reversed in TG-aKO mice. MAGL expression is elevated in astrocytes from both AD patients and TG mice. Remarkably, a single intracerebroventricular (ICV) injection of AAV-MAGL-shRNA administered at either presymptomatic or postsymptomatic stages prevents or reverses neuropathology and synaptic and cognitive impairments in TG mice, providing preclinical evidence that astrocytic MAGL silencing represents an effective therapeutic strategy for AD.

## Introduction

Alzheimer’s disease (AD) is the most common cause of dementia in the elderly. Currently, more than 6.9 million people in the United States and 55 million people worldwide are living with AD related dementia (ARAD); these numbers are projected to rise to 12.7 million in the United States and 153 million globally by 2050. Despite decades of intense effort, no therapies effectively prevent, treat, or halt AD progression. While continued elucidation of disease mechanisms is essential, there is an urgent public health need to develop novel, effective interventions.

The etiology and progression of AD are now recognized as multifactorial, involving diverse mechanisms and signaling pathways. In this context, the endocannabinoid system has emerged as a promising therapeutic avenue for AD ^1,2^, given its broad roles in physiological, pharmacological, and pathological processes ^3,4^. Endocannabinoids are endogenous lipid signaling mediators, distinct from exogenous cannabinoids such as Δ^9^-tetrahydrocannabinol (Δ^9^-THC), the principal psychoactive constituent of cannabis. Among these, 2-arachidonoylglycerol (2-AG) is the most abundant endocannabinoid. Accumulating evidence indicates that 2-AG plays a critical role in maintaining brain homeostasis by modulating synaptic transmission and plasticity, resolving neuroinflammation, and protecting neurons from harmful insults ^1,5–13^. Thus, enhancing 2-AG signaling is broadly beneficial in the brain.

Following synthesis, however, 2-AG is rapidly degraded by several enzymes, with monoacylglycerol lipase (MAGL) serving as the principal hydrolase in the brain ^1,14–17^. While 2-AG exerts anti-inflammatory and neuroprotective effects, its hydrolysis product, arachidonic acid, serves as a precursor for pro-inflammatory and neurotoxic prostaglandins (*e.g.*, PGE2) and leukotrienes (*e.g.*, LTB4) ^18–20^. Therefore, MAGL inhibition confers a “*dual effect*”: it augments beneficial 2-AG signaling while concurrently reducing pro-inflammatory, neurotoxic eicosanoids ^1,5,13,16,21–24^. Consistent with this mechanism, early studies demonstrated that systemic MAGL inactivation alleviates neuropathology and improves synaptic function and cognitive performance in AD animal models ^23–28^, supporting MAGL as a potential therapeutic target for AD ^1,23,25,29–32^.

However, accumulating evidence suggests that systemic MAGL inactivation, whether pharmacological inhibition or genetic deletion, can elicit adverse effects in both the brain and peripheral tissues ^33–39^. Importantly, selective inactivation of MAGL in neurons alone was recently shown to impair synaptic and cognitive functions ^40^, suggesting that intact neuronal 2-AG metabolism is critical for maintaining normal cognition. Because MAGL also catalyzes the final step of triglyceride lipolysis, its systemic inactivation may potentially alter lipid and energy homeostasis ^41^. Together, these findings indicate that global MAGL inactivation may not achieve the optimal therapeutic outcome for AD.

In line with this view, we recently found that astrocyte-specific MAGL deletion, but not neuron-specific inactivation, alleviates neuropathology, reduces neuroinflammation, and improves synaptic and cognitive functions in a traumatic brain injury (TBI) model ^42^. These results suggest that the neuroprotection observed with systemic MAGL inactivation arises primarily from limiting 2-AG degradation in astrocytes rather than in neurons, thereby revealing a cell type-specific role for 2-AG signaling in both physiology and disease ^1,42,43^. Accordingly, we pursued a cell type-specific strategy to target MAGL in AD.

## Results

### Astrocytic MAGL deletion prevents cognitive decline in 5xFAD mice

To delineate cell type-specific roles of 2-AG metabolism in AD, we crossed 5xFAD transgenic (TG) mice with *mgll* (encoding MAGL) lines, including wild-type *mgll*^flox/flox^ (naïve), global knockout (tKO), astrocytic knockout (aKO), and neuronal knockout (nKO) ^42^. This strategy generated 5xFAD TG cohorts with global (TG-tKO), astrocyte-specific (TG-aKO), or neuron-specific (TG-nKO) deletion of *mgll*, alongside a non-transgenic, non-knockout controls (WT). Because microglial MAGL contributes minimally to brain 2-AG hydrolysis ^17^, we did not generate 5xFAD mice lacking MAGL in microglia ^42^.

Given that cognitive impairment is the hallmark of AD, we first assessed spatial learning and memory in 6-month-old mice using the classic Morris water maze (MWM) and novel object recognition (NOR) paradigms, as 5xFAD TG mice typically begin to exhibit cognitive deficits around this age ^25,27^. As shown in Figure 1A, TG mice displayed impaired learning and memory in the MWM. This deficit was significantly attenuated in both TG-tKO and TG-aKO mice; notably, TG-aKO performance was preserved at WT levels and even exceeded WT on the probe trial assessing memory retention (Figure 1B). In contrast, TG-nKO mice exhibited deficits comparable to TG mice. Consistently, NOR testing confirmed these findings: TG-nKO mice showed impaired recognition memory similar to TG mice, TG-tKO and TG-aKO mice exhibited no such deficits (Figure 1C). Together, these findings indicate that inhibition of 2-AG degradation in astrocytes, but not in neurons, prevents cognitive decline in AD mice. The cognitive improvement observed in TG-tKO animals likely results from suppression of 2-AG metabolism in astrocytes.

**Figure 1.**
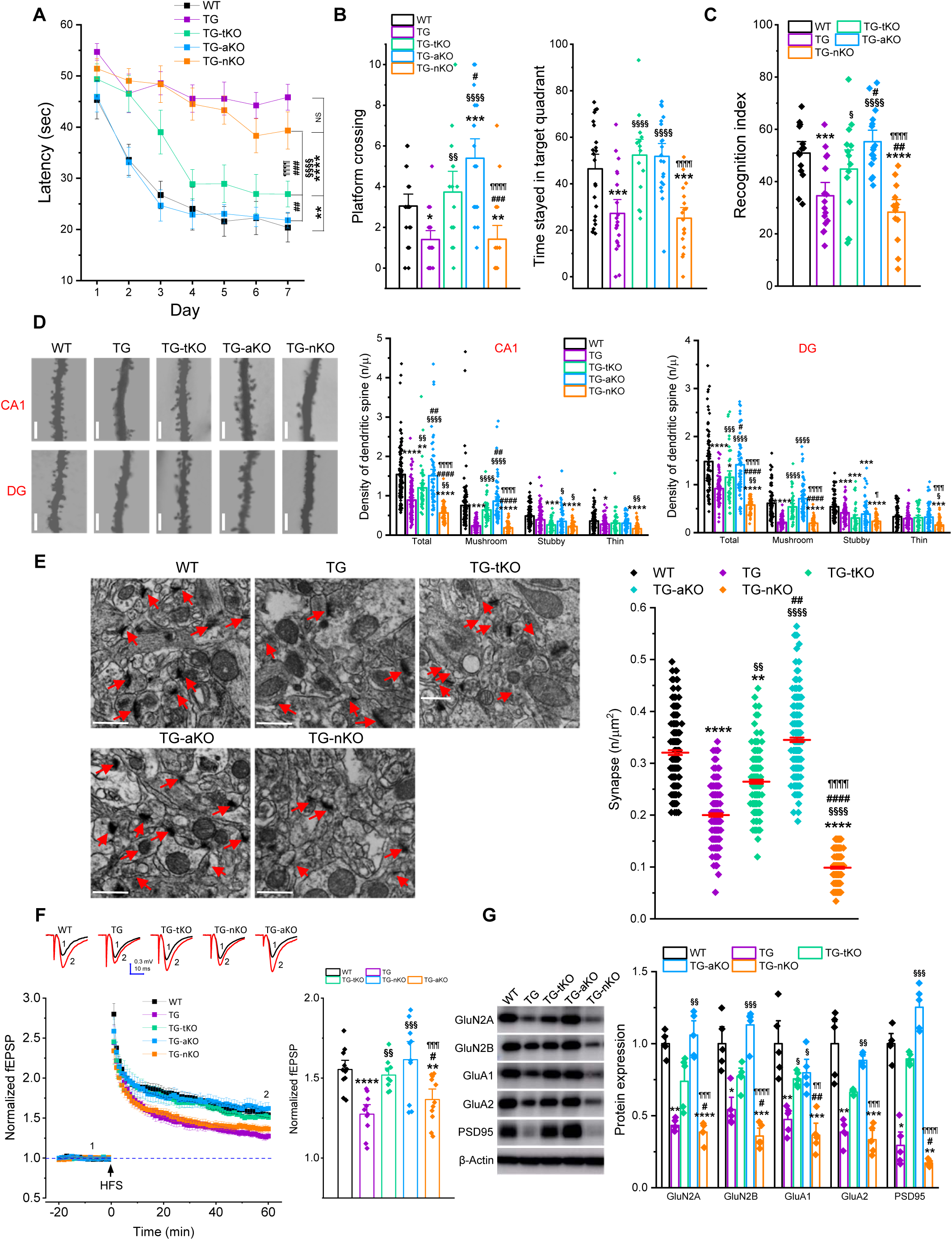
Inactivation of astrocytic MAGL prevents synaptic and cognitive deterioration in 5xFAD TG mice. (A) Spatial learning and memory were assessed using the Morris water maze (MWM) test in 6-month-old WT, TG, TG-tKO, TG-aKO, and TG-nKO mice. Data are presented as mean ± SEM. **P<0.01, ****P<0.0001 vs WT; ^§§§§^P<0.0001 vs TG; ^###^P<0.001 vs TG-tKO; ^¶¶¶¶^P<0.0001 compared with TG-aKO (Repeated measures ANOVA, n=15∼20 animals/group). (B) The probe test was conducted 24 hours following 7 days of hidden platform training. Data are presented as mean ± SEM. *P<0.05, **P<0.01, ***P<0.001 vs WT; ^§§^P<0.01, ^§§§^P<0.001 vs TG; ^#^P<0.05. ^##^P<0.01 vs TG-tKO; ^¶¶¶¶^P<0.0001 vs TG-aKO (ANOVA with Bonferroni post-hoc test). (C) Novel object recognition (NOR) test in WT and TG cohorts with *mgll* deletion. Data are presented as mean ± SEM. ***P<0.001, ****P<0.0001 vs WT; ^§^P<0.01, ^§§§§^P<0.0001 vs TG; ^#^P<0.05. ^##^P<0.001 vs TG-tKO; ^¶¶¶¶^P<0.0001 vs TG-aKO (ANOVA with Bonferroni post-hoc test, n=13∼16 animals/group). (D) Golgi staining of dendritic spines in hippocampal CA1 pyramidal neurons and dentate gyrus granule neurons. Data are presented as mean ± SEM. *P<0.05, **P<0.01, ***P<0.001, ****P<0.0001 vs WT; ^§^P<0.05, ^§§^P<0.01, ^§§§^P<0.001, ^§§§§^P<0.0001 vs TG; ^#^P<0.05. ^##^P<0.01, ^####^P<0.01 vs TG-tKO; ^¶^P<0.05, ^¶¶¶^P<0.001, ^¶¶¶¶^P<0.0001 vs TG-aKO (ANOVA with Bonferroni post-hoc test). (E) Transmission electron microscopy (TEM) images of hippocampal sections from WT, TG, TG-tKO, TG-aKO, and TG-nKO mice. Data are mean ± SEM. **P<0.01, ****P<0.0001 vs WT; ^§§^P<0.01, ^§§§§^P<0.0001 vs TG; ^##^P<0.01, ^####^P<0.01 vs TG-tKO; ^¶¶¶¶^P<0.0001 vs TG-aKO (ANOVA with Bonferroni post-hoc test, n=5 animals/group). (F) Lon-term potentiation (LTP) recorded at hippocampal CA3-CA1 synapses. Data are presented as mean ± SEM. **P<0.01, ****P<0.0001 vs WT; ^§§^P<0.01, ^§§§^P<0.001 vs TG; ^#^P<0.05 vs TG-tKO; ^¶¶¶^P<0.001 vs TG-aKO (ANOVA with Bonferroni post-hoc test, n=5∼6 animals/group). (G) Immunoblot analysis of hippocampal expression of glutamate receptor subunits and postsynaptic density protein 95 (PSD-95). Data are presented as mean ± SEM. **P<0.01, ***P<0.001, ****P<0.0001 vs; ^§^P<0.05, ^§§^P<0.01, ^§§§^P<0.001 vs TG; ^#^P<0.05, ^##^P<0.01 vs TG-tKO; ^¶¶¶^P<0.001 vs TG-aKO (Kruskal-Wallis test with Dunn’s post hoc test for multiple comparisons, n=5 animals/group).

### Astrocytic MAGL inhibition preserves synaptic structure and function in AD mice

Because cognitive function depends on intact synaptic structure and function, we assessed hippocampal synapses in the mouse cohorts shown in Figure 1A-C. Dendritic spines, specialized postsynaptic protrusions on dendrites, were significantly reduced in pyramidal and granule neurons in both the CA1 and dentate gyrus (DG) regions of TG mice, but preserved in TG-tKO and TG-aKO mice (Figure 1D). By contrast, TG-nKO mice exhibited a further reduction in spine density, lower than in TG mice. Consistent with this, transmission electron microscopy (TEM), which allows direct visualization of individual synapses, revealed a marked reduction in hippocampal synapse number in TG mice. This reduction was attenuated in TG-tKO mice, while synapse density in TG-aKO mice was fully restored to WT levels. By contrast, TG-nKO mice exhibited an even greater loss of synapses compared with WT (Figure 1E).

We next evaluated long-term potentiation (LTP), a cellular model of learning and memory, at hippocampal CA3–CA1 synapses and found that LTP was impaired in TG mice but remained normal in TG-tKO and TG-aKO mice, whereas TG-nKO mice exhibited impairments comparable to TG mice (Figure 1F). The impaired LTP in TG and TG-nKO mice likely reflects reduced glutamate receptor expression. Indeed, as shown in Figure 1G, levels of AMPA and NMDA receptor subunits, as well as the synaptic marker PSD95, were significantly decreased in TG and TG-nKO mice, but not in TG-tKO or TG-aKO mice. Consistently, Aβ treatment of mixed neuron-glia cultures from WT, tKO, aKO, and nKO mice revealed that astrocytic MAGL deletion (aKO) confers resilience to Aβ-induced reductions in synaptic markers, including PSD95 and VGluT1, whereas this protection was not observed in nKO cultures (Supplementary Figure 1).

Together, these results demonstrate that astrocyte-specific MAGL inactivation preserves hippocampal synaptic structure and function in AD mice, whereas neuronal MAGL inactivation fails to confer protection and instead worsens synaptic deterioration.

### Astrocytic MAGL inactivation ameliorates Aβ neuropathology in AD mice

Extracellular Aβ plaque deposition is a central neuropathological hallmark of AD. To determine whether astrocyte-specific inactivation of MAGL attenuates Aβ pathology, we assessed Aβ burden in TG mice and in TG mice lacking MAGL. As shown in Figure 2A, TG mice exhibited a high burden of 4G8-immunopositive Aβ plaques, which was markedly reduced in TG-tKO and TG-aKO mice, but showed little to no reduction in TG-nKO mice. Consistently, soluble Aβ42 levels in hippocampal tissue were significantly decreased in TG-tKO and TG-aKO mice compared with TG mice, whereas TG-nKO mice showed no reduction (Figure 2B). These findings suggest that the alleviation of Aβ pathology observed in TG-tKO and TG-aKO mice likely reflects the restriction of 2-AG breakdown in astrocytes.

**Figure 2.**
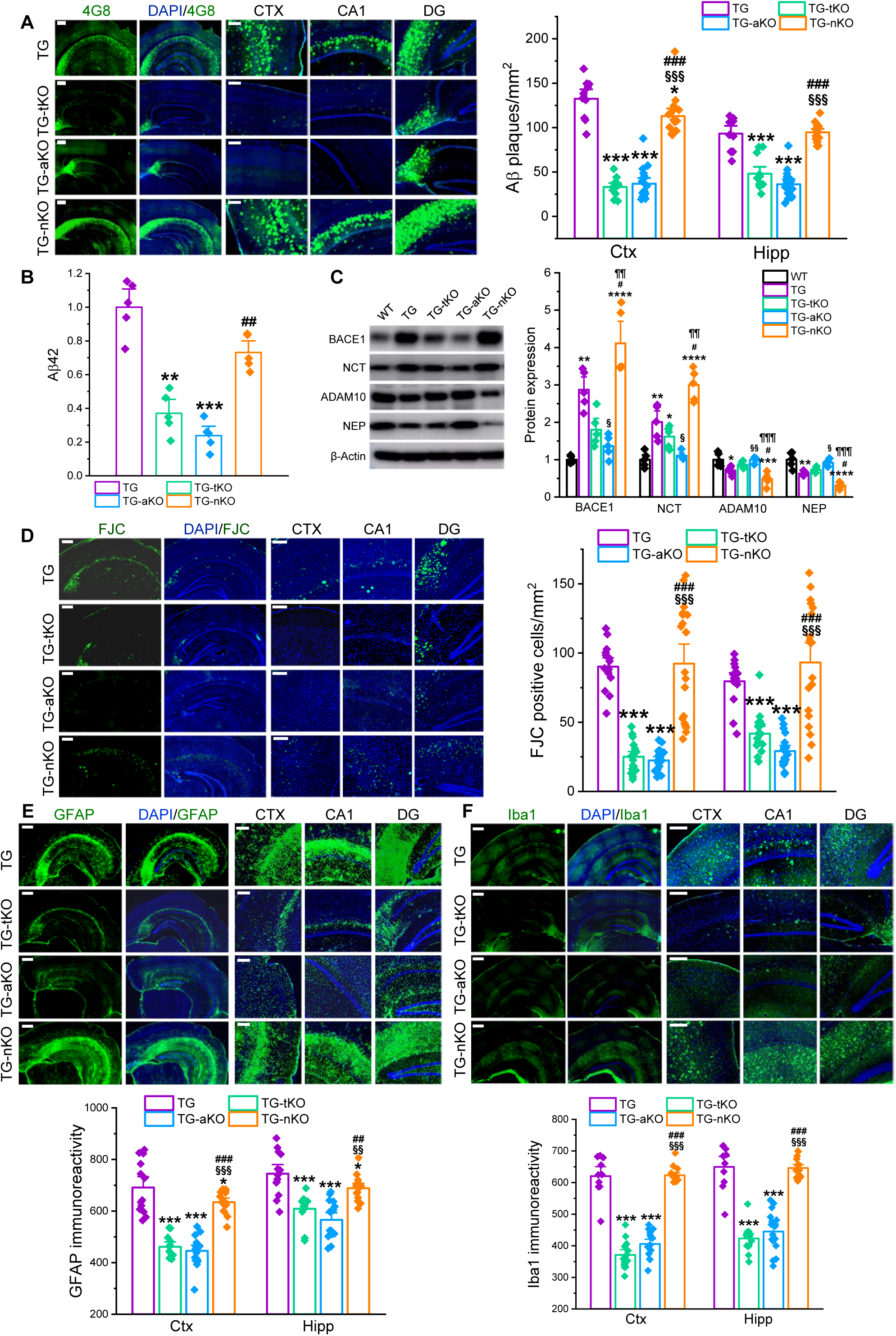
Astrocytic MAGL inactivation attenuates neuroinflammation, Aβ pathology, and neurodegeneration in TG mice. (A) Aβ plaques, assesses by 4G8 immunoreactivity, are reduced in the brains of TG-tKO and TG-aKO mice, but no in TG-nKO mice. Data are presented as mean ± SEM. *P<0.05, ***P<0.001 vs TG; ^§§§^P<0.001 vs TG-tKO; ^###^P<0.001 vs TG-aKO (ANOVA with Bonferroni post-hoc test, n=5 animals/group). Scale bars: 400 µm. (B) Hippocampal soluble Aβ42 levels assessed by ELISA are reduced in both TG-tKO and TG-aKO mice, but no in TG-nKO mice. Data are presented as mean ± SEM. **P<0.01, ***P<0.001 vs TG; ^##^P<0.01 vs TG-aKO (Kruskal-Wallis test with Dunn’s post hoc test for multiple comparisons, n=5 animals/group). (C) Immunoblot analysis of Aβ-processing enzymes in the hippocampus of WT, TG, TG-tKO, TG-aKO, and TG-nKO mice. Data are presented as mean ± SEM. *P<0.05, **P<0.01, ***P<0.001, ****P<0.0001 vs WT; ^§^P<0.05, ^§§^P<0.01 vs TG; #P<0.05 vs TG-tKO; ^¶¶^P<0.01, ^¶¶¶¶^P<0.0001 vs TG-aKO (Kruskal-Wallis test with Dunn’s post hoc test for multiple comparisons, n=5 animals/group). (D) FJC staining of degenerating neurons in the brain. Data are mean ± SEM. ***P<0.001 vs TG; ^§§§^P<0.001 vs TG-tKO; ^###^P<0.001 vs TG-aKO (ANOVA with Bonferroni post-hoc test, n=5 animals/group). Scale bars: 200 µm. (E∼F) Immunostaining of astrocytic and microglial reactivity. Data are presented as mean ± SEM. *P<0.05, ***P<0.001 vs TG; ^§§^P<0.01, ^§§§^P<0.001 vs TG-tKO; ^##^P<0.01, ^###^P<0.001 vs TG-aKO (ANOVA with Bonferroni post-hoc test, n=5 animals/group). Scale bars: 400 µm.

Aβ production depends on multiple processing enzymes. Accordingly, we assessed hippocampal expression of BACE1 (β-secretase), nicastrin (NCT, a core component of the γ-secretase complex), ADAM10 (α-secretase), and neprilysin (NEP, an Aβ-degrading enzyme). As shown in Figure 2C, BACE1 and NCT were significantly elevated, whereas ADAM10 and NEP were reduced in TG mice. These changes were attenuated or reversed in TG-tKO and TG-aKO mice, with more prominent effects in TG-aKO mice. In contrast, expressions of these enzymes in TG-nKO mice were comparable to, or worse than, that in TG mice. These data indicate that astrocytic MAGL inactivation suppresses Aβ-producing enzymes while enhancing Aβ-degrading enzymes, suggesting that elevated astrocytic 2-AG signaling shifts both secretase and clearance pathways, potentially underlying the reduced plaque burden and Aβ42 levels observed in TG-tKO and TG-aKO mice.

### Astrocytic MAGL inactivation reduces neurodegeneration in AD mice

Neurodegeneration is a defining feature of AD. We quantified degenerating neurons in TG mice and TG mice lacking MAGL using Fluoro-Jade C (FJC), a marker of degenerating neurons. As shown in Figure 2D, FJC-positive neurons were significantly reduced in TG-tKO and TG-aKO mice compared with TG mice, but not in TG-nKO mice, indicating that astrocytic, rather than neuronal, MAGL inactivation confers neuroprotection in AD, suggesting astrocytic 2-AG signaling mediates astrocyte-neuron interactions that limit neurodegeneration.

### Astrocytic MAGL inactivation reduces neuroinflammation in AD mice

Neuroinflammation is a key driver of neurodegenerative diseases ^44–46^. Therefore, we assessed astrocytic and microglial reactivity as readouts of neuroinflammation and found that GFAP and Iba1 immunoreactivity were significantly attenuated in TG-tKO and TG-aKO mice when compared with TG mice, but not in TG-nKO mice (Figure 2E-F). These results indicate that inhibiting 2-AG degradation in astrocytes alleviates neuroinflammation in AD mice, suggesting that astrocytic 2-AG acts through autocrine signaling and astrocyte-microglia interactions to resolve neuroinflammation.

### *Astrocytic MAGL inactivation* reverses dysregulated expression of synaptic, inflammatory, and AD-related genes in AD mice

As shown above, selective MAGL inactivation in astrocytes, but not neurons, prevents cognitive decline, preserves synaptic structure and function, and reduces Aβ pathology, neurodegeneration, and neuroinflammation in TG mice. To probe underlying cellular and molecular mechanisms, we profiled hippocampal transcriptomes by single-nucleus RNA-seq (snRNA-seq), as described previously ^42,47^. We observed that numerous genes were differentially expressed across genotypes: dysregulated genes exhibited in TG mice were significantly attenuated or reversed in TG-tKO and TG-aKO mice, approaching to WT levels, but not in TG-nKO mice (Figure 3A and Supplementary Table S1). Functional annotations via GO enrichment analysis showed changes in differentially expressed genes (DEGs) involving presynaptic endocytosis, and NF-kB signaling pathways in TG-aKO mice were opposite from TG mice, suggesting that altered functions were significantly reversed by astrocyte-specific inactivation of MAGL in TG mice (Supplementary Table 2). Among the top dysregulated genes implicated in AD and synaptic function (*e.g.*, *cntn2*, *mid1*, *tyro3*, *fmnl2*, and *dlg3*) ^48–52^, expression abnormalities present in TG mice were mitigated in TG-tKO and TG-aKO, but persisted in TG-nKO mice (Figure 3B).

**Figure 3.**
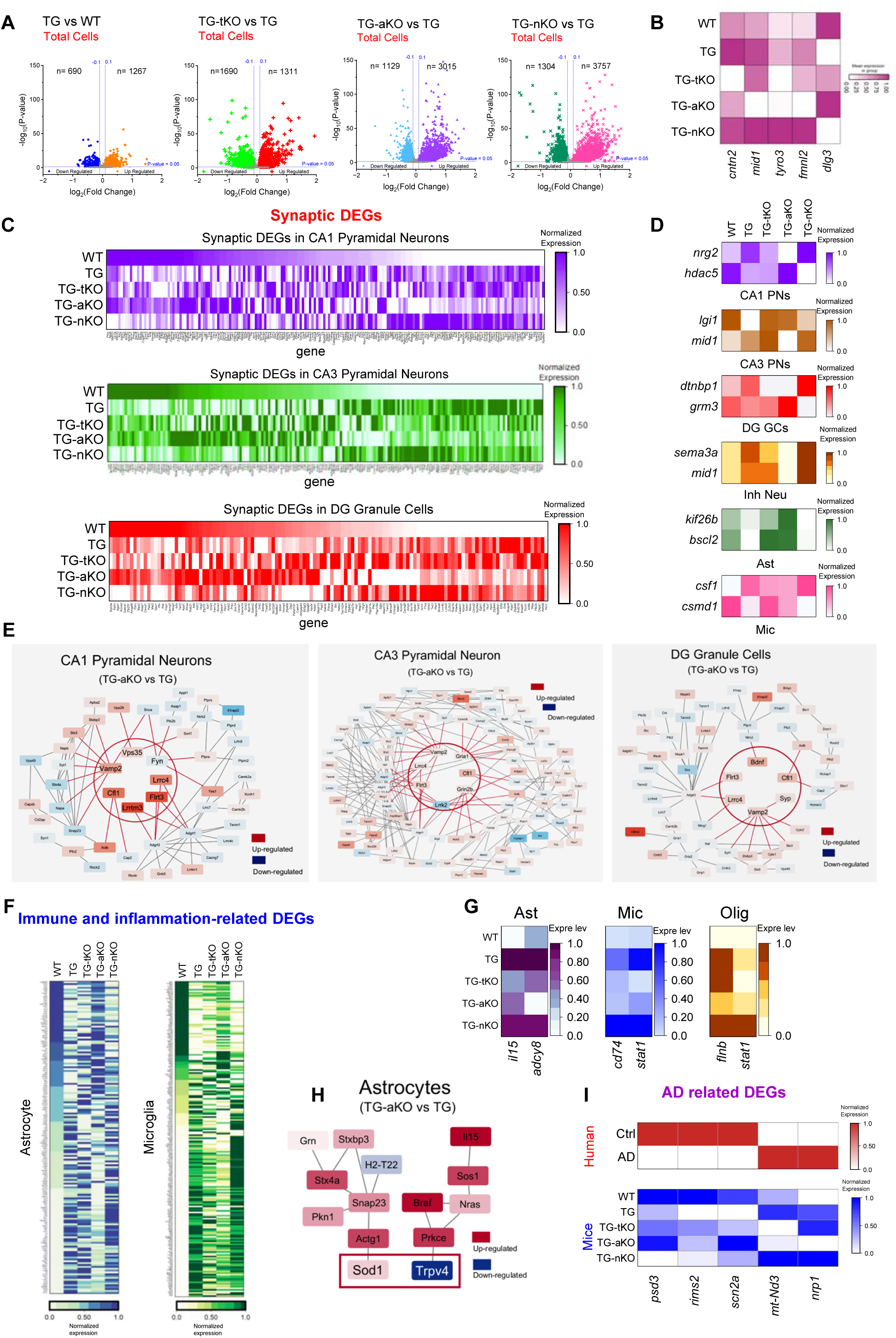
Transcriptomics analysis of synapse- and immune/inflammation-related differentially expressed genes (DEGs) in 5xFAD cohorts with *mgll* deletion. (A) Volcano plots displaying DEGs of total cells from each group. (B) Heatmap showing the top dysregulated genes implicated in AD and synaptic function. (C) Heatmaps illustrating synaptic DEGs in CA1 pyramidal neurons (PNs), CA3 PNs, and dentate gyrus granule cells (DG GCs) across groups. (D) Heatmaps showing the expression levels of representative synaptic DEGs in individual CA1 PNs, CA3 PNs, DG GCs, inhibitory neurons (Inh Neu), astrocytes (Ast), and microglia (Mic) across different transgenic mice. (E) Gene network analyses of synaptic DEGs in CA1 PNs, CA3 PNs, and DG GCs revealing potential central regulators of synaptic DEGs in TG-aKO mice. (F) Heatmaps showing expression levels of representative synaptic DEGs in Mic and Ast from different transgenic mice. (G) Heatmaps illustrating expression levels of immune/inflammation-related DEGs in Ast, Mic, and oligodendrocytes (Olig) from different groups. (H) Gene network analysis revels *sod1* and *trpv4* as key regulators of immune/inflammation-related in astrocytes of TG-aKO mice. (I) Astrocytic MAGL inactivation reverses multiple AD-related DEGs in CA1 PNs associated with synaptic function and AD pathology that are altered in AD patients and TG mice.

Next, we analyzed the expression profiles of synapse-related DEGs in hippocampal neurons and glial cells based on cell-type annotations using established markers (Supplementary Figure 2A). Expression levels of these DEGs in neurons and glial cells are presented in Supplementary Table 3 and Supplementary Figure 2B∼H. Interestingly, the expression profiles of DEGs in TG-aKO mice closely resembled those in WT mice. Likewise, synaptic DEGs in neurons and glial cells from the TG-aKO group displayed expression patterns highly similar to WT, but distinct from those in TG and TG-nKO mice (Figure 1C, Supplementary Figure 3A, and Supplementary Table 4). We identified several DEGs involved in synaptic transmission, synapse formation, and synapse maturation ^52–61^, including *nrg2* and *hdac5* in CA1 pyramidal neurons (PNs); *lgi1* and *mid1* in CA3 PNs; *dtnbp1* and *grm3* in dentate gyrus granule cells (DGGCs); *sema3a* and *mid1* in interneurons (Inh Neu); *kif26b* and *bscl2* in astrocytes (Ast); and *csf1* and *csmd1* in microglia (Mic). These genes were dysregulated in TG mice but were normalized or shifted toward WT levels in TG-tKO and TG-aKO mice, more prominently in TG-aKO mice, whereas their expression in TG-nKO mice remained similar to that in TG mice (Figure 3D). To investigate how synaptic DEGs are regulated in TG-aKO mice to counteract the alterations observed in TG mice, we constructed a gene network of synaptic DEGs (Figure 3E). This analysis revealed that *cfl1*, *flrt3*, *lrrc4*, and *vamp2* function as shared central regulators, whereas *fyn*, *lrrtm3*, *vps35*, *gria1*, *lrrk2*, *grin2b*, *bdnf*, and *syp* act as cell type-specific regulators in TG-aKO mice.

We then analyzed immune/inflammation-related DEGs and found that expression levels in TG-aKO mice were nearly identical to those in WT mice, whereas these DEGs in TG-nKO mice resembled those in TG mice (Figure 1F, Supplementary Figure 2B, and Supplementary Table 5). Specifically, several immune/inflammation-related DEGs that were upregulated in TG mice, including *il15* and *adcy8* in Ast, *cd74* and *stat1* in Mic, and *flnb* and *stat1* in Olig ^62–67^, were significantly downregulated in TG-aKO mice to levels comparable to WT mice (Figure 1G), but not in TG-nKO mice. Gene network analysis indicates that *sod1* and *trpv4* might be key regulators of these DEGs in astrocytes of TG-aKO mice (Figure 1H).

Finally, we analyzed the effects of inhibiting astrocytic 2-AG degradation on AD-related DEGs in TG mice. As shown in Supplementary Figure 3C and Supplementary Table 6, many AD-related DEGs across different cell types in WT and TG-aKO mice exhibited similar expression patterns, whereas they were altered in the opposite direction in TG and TG-nKO mice. To determine whether AD-related DEGs in patients with AD resemble those in mice, we reanalyzed published snRNA-seq data from the human hippocampus and re-annotated the clusters (Supplementary Figure 3D&E). As shown in Figure 3I, multiple AD-related DEGs in CA1 PNs, such as *psd3*, *rims2*, *scn2a*, *mt-ND3*, and *nrp1*, associated with synaptic function and AD pathology, showed similar changes in AD patients and TG mice ^68–72^. Importantly, these dysregulated DEGs were restored to WT levels in both TG-tKO and TG-aKO mice, albeit to a slightly lesser extent in TG-tKO mice, but were not corrected in TG-nKO mice. Together, our transcriptomic data suggest that inhibition of 2-AG degradation in astrocytes in TG-aKO mice prevents or reverses aberrant expression of many synapse-, immune/inflammation-, and AD-related DEGs, which may contribute to attenuated neuropathology and improved synaptic and cognitive functions in TG mice.

### Intrahippocampal silencing of MAGL in astrocytes prevents synaptic and cognitive decline in TG mice

Our findings from TG mice crossed with MAGL knockout lines demonstrate that inhibiting astrocytic 2-AG degradation plays a crucial role in mitigating neuropathology and preserving synaptic and cognitive function in AD animals. Based on this, we hypothesized that silencing MAGL in astrocytes would alleviate synaptic and cognitive deficits in TG mice. To test this, AAV5 vectors driven by the GFAP promoter and expressing either *mgll*-shRNA or a control construct were stereotaxically injected into the hippocampus of 4-month-old TG mice. Synaptic and cognitive functions were assessed at 6 months of age, 2 months after AAV injection (Figure 4A, Supplementary Figure 4A&B). TG mice receiving control AAV exhibited impaired learning and memory in both NOR and MWM, whereas AAV-mgll-shRNA-treated TG mice showed improved behavioral performance (Figure 4B∼D). Moreover, astrocyte-selective MAGL knockdown prevented LTP impairment (Figure 4E) and restored the expression of glutamate receptor subunits and the synaptic marker PSD95, which were reduced in TG mice (Figure 4F). Together, these results indicate that silencing hippocampal astrocytic MAGL, thereby limiting 2-AG breakdown, prevents synaptic and cognitive deterioration in TG mice.

**Figure 4.**
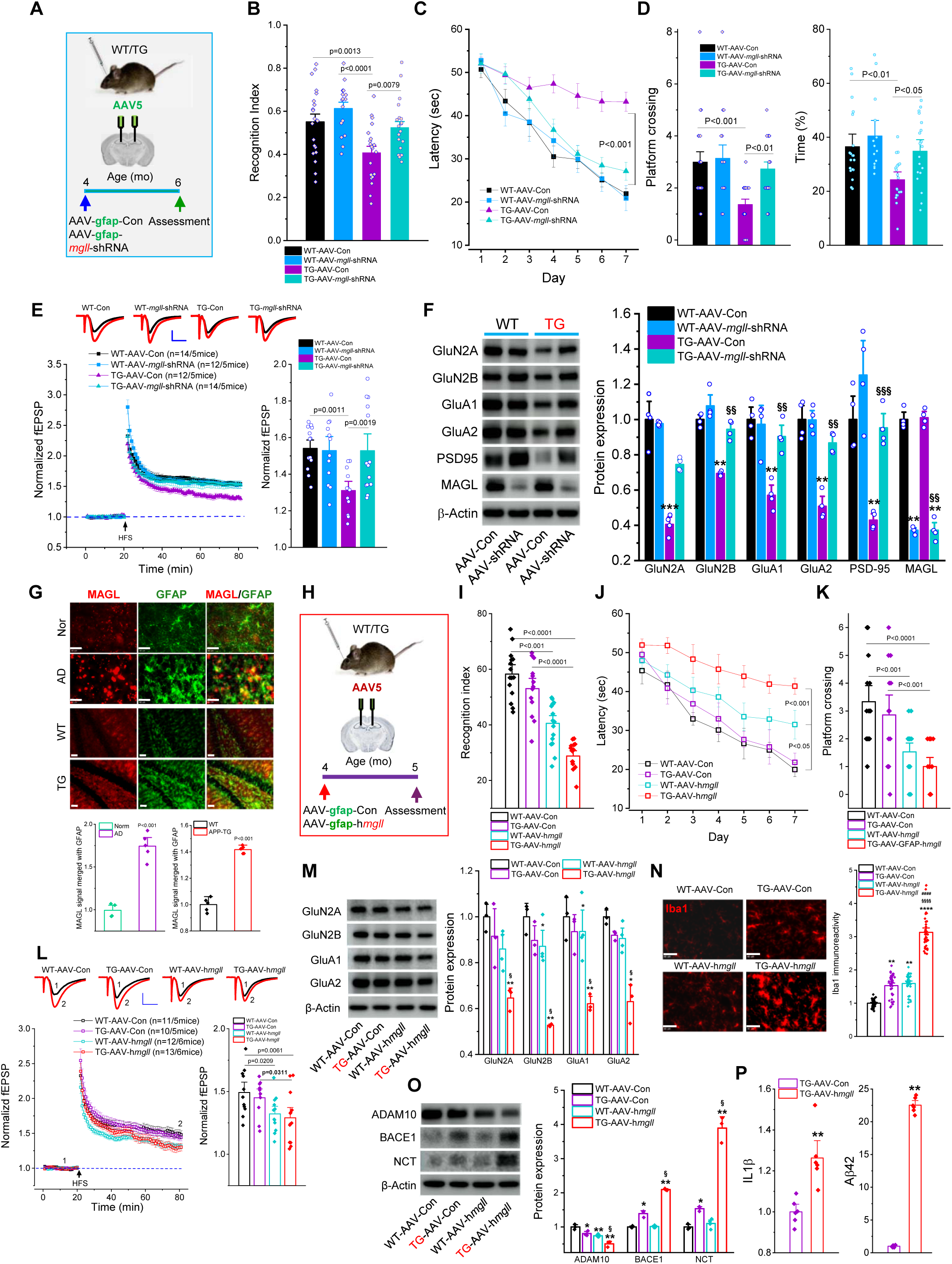
Silencing astrocytic MAGL produces protection, whereas its overexpressing exacerbates neuropathology and accelerates synaptic and cognitive decline in TG mice. (A) Schematic illustration of the protocol for AAV-*mgll*-shRNA injection. (B) NOR test in WT and TG mice injected with AAV-Con or AAV-mgll-shRNA. Data presented as mean ± SEM (ANOVA with Bonferroni post-hoc test, n=16∼20 animals/group). (C) Learning acquisition in the MWM test. The data are presented as mean ± SEM (Repeated measures ANOVA, n=14∼19 animals/group). (D) Probe test performed 24 hours after 7 days of hidden platform training. Data are presented as mean ± SEM (ANOVA with Bonferroni post-hoc test). (E) LTP at hippocampal CA3-CA1 synapses recorded from WT and TG mice treated with AAV-Con or AAV-*mgll*-shRNA. Data are presented as mean ± SEM (ANOVA with Bonferroni post-hoc test, n=5 animals/group). (F) Immunoblot analysis of hippocampal expression of glutamate receptor subunits and PSD-95. Data are presented as mean ± SEM. **P<0.01, ***P<0.001 vs WT-AAV-Con, ^§§^P<0.01, ^§§§^P<0.001 vs TG-AAV-Con (Kruskal-Wallis test with Dunn’s post hoc test for multiple comparisons, n=4 animals/group). (G) Immunostaining of MAGL and GFAP in the hippocampus of AD patients and TG mice. Data are presented as mean ± SEM (Student’s t-test, n=5 subjects/animals/group). Scale bars: 20 µm. (H) Schematic illustration of the protocol for AAV-h*mgll* overexpression. (I) NOR test in WT and TG mice injected with AAV-Con or AAV-h*mgll*. Data presented as mean ± SEM (ANOVA with Bonferroni post-hoc test, n=14∼17 animals/group). (J) Learning acquisition in the MWM test. Data are presented as mean ± SEM (Repeated measures ANOVA). (K) The probe test. The data are presented as mean ± SEM (ANOVA with Bonferroni post-hoc test, n=14∼17 animals/group). (L) LTP at hippocampal CA3-CA1 synapses recorded from WT and TG mice injected with AAV-Con or AAV-h*mgll*. Data are presented as mean ± SEM (ANOVA with Bonferroni post-hoc test, n=5∼6 animals/group). (M) Immunoblot analysis of hippocampal expression of glutamate receptor subunits. Data are presented as mean ± SEM. *P<0.05, **P<0.01 vs WT-AAV-Con, ^§^P<0.05 vs TG-AAV-Con (Kruskal-Wallis test with Dunn’s post hoc test for multiple comparisons, n=4∼5 animals/group). (N) Immunostaining of microglial reactivity in the hippocampus of the animals overexpressing human MAGL. Data are presented as mean ± SEM (ANOVA with Bonferroni post-hoc test, n=5 animals/group). Scale bars: 40 µm. (O) Immunoblot analysis of hippocampal expression of Aβ-processing enzymes. Data are presented as mean ± SEM. *P<0.05, **P<0.01 vs WT-AAV-Con, ^§^P<0.05 vs TG-AAV-Con (Kruskal-Wallis test with Dunn’s post hoc test for multiple comparisons, n=3∼4 animals/group). (P) ELISA analysis of hippocampal expression of IL-1β and Aβ42 levels. Data are presented as mean ± SEM. **P<0.01 vs TG-AAV-Con (Mann-Whitney test, n=6 animals/group).

### Astrocytic MAGL overexpression accelerates synaptic and cognitive decline in TG mice

Prior studies have reported elevated MAGL expression or activity in the brains of individuals with AD and in AD model animals ^73–75^, but it remains unclear whether MAGL is specifically upregulated in astrocytes. Using double-label immunostaining, we observed elevated expression of MAGL in the hippocampus of postmortem AD brains and in 6-month-old TG mice (Figure 4G). However, direct evidence linking astrocyte-specific MAGL upregulation to synaptic and cognitive deficits has been lacking. To address this, we used the same AAV5 vector for *mgll*-shRNA driven by a *GFAP* promoter to overexpress human *mgll* selectively in astrocytes (Figure 4H, Supplementary Figure-4C&D). TG mice receiving control AAV showed no significant learning or memory deficits at 5 months of age, whereas astrocyte-specific MAGL overexpression markedly impaired performance on both NOR and MWM tests (Figure 4I-K). Notably, astrocytic MAGL overexpression in WT mice also impaired behavioral performance. Impaired in LTP and reduced expression of glutamate receptor subunits in TG and WT mice overexpressing *mgll* further support that elevated astrocytic MAGL is detrimental to synaptic and cognitive functions (Figure 4 L&M). These results indicate that increased astrocytic MAGL expression accelerates cognitive decline in presymptomatic TG mice and impairs cognition even in WT controls, providing the first evidence that astrocytic 2-AG metabolism is directly linked to cognitive function in both physiological and AD contexts.

In addition, astrocytic MAGL overexpression elicited microglial reactivity in WT mice and further exacerbated it in TG mice (Figure 4N), while reducing ADAM10 and increasing BACE1 and NCT expression in TG mice (Figure 4O). ELISA measurements showed robust increases in IL-1β and Aβ42 in TG mice receiving AAV overexpressing human *mgll* (Figure 4P), indicating that heightened astrocytic MAGL not only accelerates synaptic and cognitive decline but also exacerbates neuroinflammation and promotes Aβ production in TG mice.

### Intracerebroventricular delivery of AAV silencing astrocytic MAGL at the presymptomatic stage prevents the onset of synaptic and cognitive decline, as well as neuropathological deterioration, in TG mice

The evidence from TG mice with astrocytic *mgll* deletion, as well as AAV-mediated astrocytic *mgll* silencing or overexpression, demonstrates that astrocytic MAGL is a key pathogenic factor and viable therapeutic target in AD. However, current pharmacotherapies cannot target genes in a cell type-specific manner in the brain. Therefore, we pursued a clinically relevant approach using intracerebroventricular (ICV) delivery of AAV-*mgll*-shRNA vectors to further validate astrocytic MAGL as a therapeutic target. To enhance gene transfer efficiency, we used AAV-PHP-eB, a recently developed variant of AAV9 ^76,77^, and applied the same cloning strategy used for AAV5 to generate to generate *AAV-PHP.eB-GFAP-eGFP-mgll-shRNAmir(2)* for selective MAGL silencing in astrocytes; *AAV-PHP.eB-GFAP-eGFP* served as the control. ICV delivery of AAV-PHP.eB-*mgll*-shRNA with a gfap promoter transduced primarily astrocytes, with minimal or no neuronal transduction (Supplementary Figure 5A&B), and produced a robust reduction of MAGL expression in astrocytes (Supplementary Figure 5C).

To determine whether silencing astrocytic MAGL prevents the onset of synaptic and cognitive decline in TG mice, AAV-*mgll*-shRNA was delivered at 4 months of age (presymptomatic), and assessments were performed at 6 months of age, 2 months post-injection (Figure 5A). We first evaluated cognitive function using the MWM and NOR tests. As shown in Figure 5B∼D, TG mice treated with control AAV displayed deficits in learning and memory compared with WT mice, whereas these deficits were prevented in TG mice receiving AAV-*mgll*-shRNA. The NOR test also showed improved memory retention in AAV-*mgll*-shRNA-treated TG mice (Figure 5E). Consistently, ICV delivery of AAV-*mgll*-shRNA prevented synaptic deterioration in TG mice: impaired LTP was rescued by astrocytic MAGL knockdown (Figure 5F), and the reduced glutamate receptor subunits and PSD95 in TG mice were restored to levels comparable to WT mice treated with AAV-Con (Figure 5G). These data indicate that silencing MAGL in astrocytes prevents synaptic and cognitive decline in AD animals. Interestingly, WT mice treated with AAV-*mgll*-shRNA exhibited improved learning acquisition, enhanced LTP, and increased PSD95 expression compared with controls (Figure 5B, F&G), suggesting that inhibiting astrocytic 2-AG degradation enhances synaptic function and learning even under physiological conditions.

**Figure 5.**
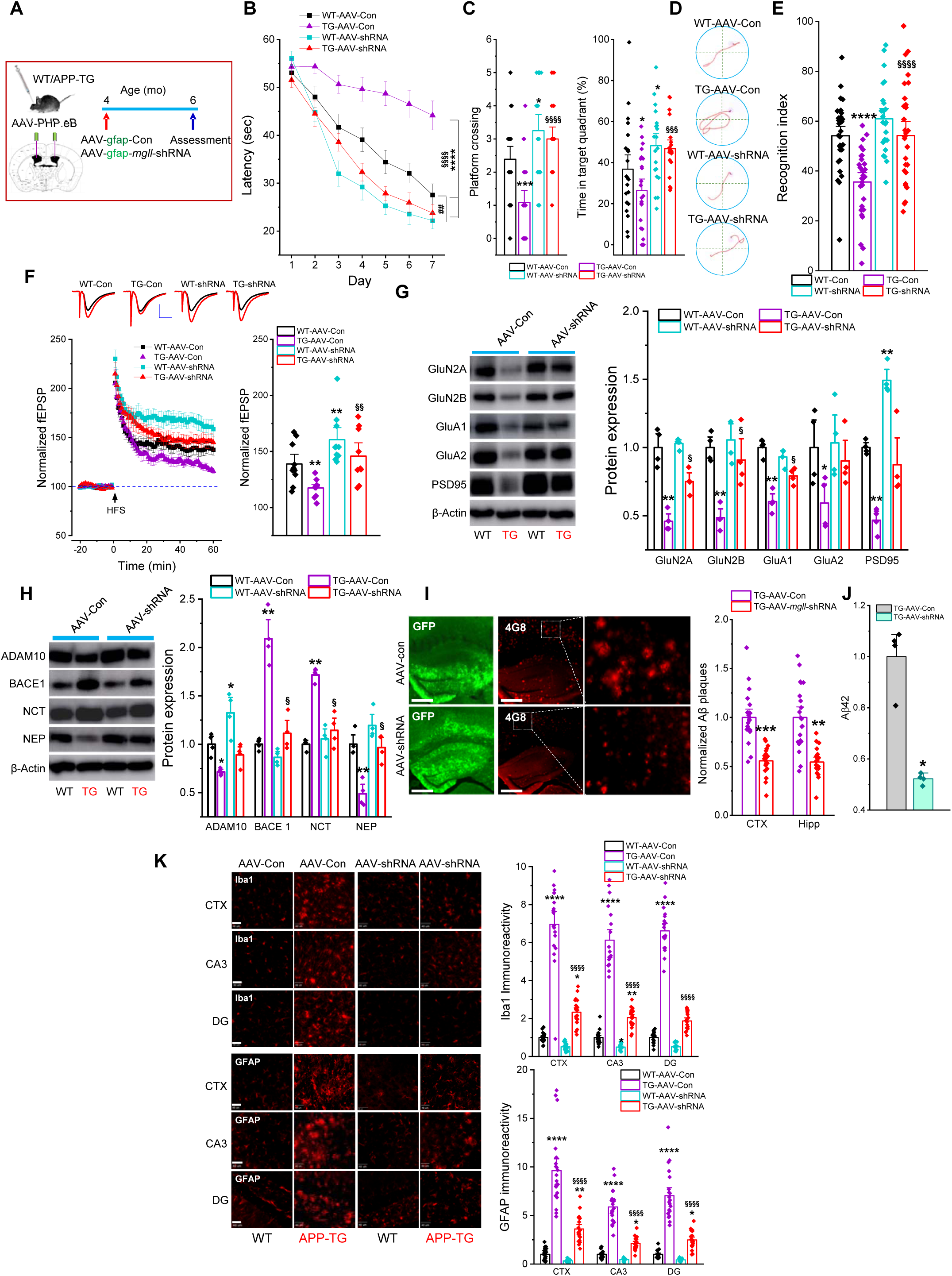
Intracerebroventricular (ICV) delivery of AAV-*mgll*-shRNA vectors at a presymptomatic age prevents the onset of neuropathology and synaptic and cognitive decline in TG mice. (A) Schematic illustration of the protocol for ICV injection of AAV-*mgll*-shRNA vectors. (B) Learning acquisition in the MWM test. Data are presented as mean ± SEM. ****P<0.0001 vs WT-AAV-Con; ^§§§§^P<0.0001 vs TG-AAV-Con; ^##^P<0.01 vs WT-AAV-shRNA (Repeated measures ANOVA). (C) Probe test. Data are presented as mean ± SEM (ANOVA with Bonferroni post-hoc test, n=20∼23 animals/group). (D) Representative swim paths during acquisition training on day 7. (E) NOR test in WT and TG mice injected with AAV-Con or AAV-*mgll*-shRNA. Data presented as mean ± SEM (ANOVA with Bonferroni post-hoc test, n=25∼30 animals/group). (F) LTP recordings at hippocampal CA3-CA1 synapses. Data are presented as mean ± SEM. **P<0.01 vs WT-AAV-Con; ^§§^P<0.01 vs TG-AAV-Con (ANOVA with Bonferroni post-hoc test, n=4∼5 animals/group). (G) Immunoblot analysis of hippocampal expression of glutamate receptor subunits and PSD95. Data are presented as mean ± SEM. *P<0.05, **P<0.01 vs WT-AAV-Con, ^§^P<0.05 vs TG-AAV-Con (Kruskal-Wallis test with Dunn’s post hoc test for multiple comparisons, n=4 animals/group). (H) Immunoblot analysis of hippocampal expression of Aβ-processing enzymes. Data are presented as mean ± SEM. *P<0.05, **P<0.01 vs WT-AAV-Con, ^§^P<0.05 vs TG-AAV-Con (Kruskal-Wallis test with Dunn’s post hoc test for multiple comparisons, n=3∼4 animals/group). (I) Immunostaining of 4G8 immunoreactivity. Data are presented as mean ± SEM. **P<0.01, ***P<0.001 vs TG-AAV-Con (ANOVA with Bonferroni post-hoc test, n=4 animals/group). Scale bars: 400 µm. (J) ELISA analysis of hippocampal Aβ42 levels. Data are presented as mean ± SEM. *P<0.05 (Mann-Whitney test, n=4 animals/group). (K) Immunostaining of GFAP and Iba1 reactivity in the brain. Data are presented as mean ± SEM. *P<0.05, **P<0.01, ****P<0.0001 vs WT-AAV-Con, ^§§§§^P<0.0001 vs TG-AAV-Con (ANOVA with Bonferroni post-hoc test, n=4 animals/group). Scale bars: 40 µm.

We next assessed neuropathology and neuroinflammation in TG mice treated with AAV-*mgll*-shRNA. Silencing astrocytes MAGL resulted in reversing altered enzymes processing Aβ, reducing Aβ and soluble Aβ42 levels, as well as attenuating reactivity of astrocytes and microglia (Figure 5H∼K).

Together, our data indicate that treatment with AAV-*mgll*-shRNA vectors at the presymptomatic stage prevents the onset of synaptic and cognitive decline, as well as deterioration in neuroinflammation and Aβ pathology, in AD animals.

### Intracerebroventricular delivery of AAV silencing astrocytic MAGL at the symptomatic stage reverses or delays the progression of synaptic and cognitive impairments, as well as neuropathological deterioration, in TG mice

To determine whether restraining astrocytic 2-AG metabolism can reverse or slow AD progression, we delivered AAV-*mgll*-shRNA vectors via ICV injection to TG mice at 6 months of age, when synaptic and cognitive deficits are already present ^25,27^, and performed assessments at 8 months of age (Figure 6A). As shown in Figure 6B∼E, impairments in learning and memory in TG mice, as measured by MWM and NOR, were reversed by astrocytic MAGL knockdown. Similarly, reductions in glutamate receptor subunits and PSD95 were rescued in TG mice by silencing astrocytic MAGL (Figure 6F). We also found that astrocytic MAGL silencing normalized the expression of Aβ-processing enzymes, reduced Aβ plaques and Aβ42 levels, and attenuated neuroinflammation (Figure 6G–J). These results suggest that inhibiting 2-AG metabolism in astrocytes can reverse or delay AD progression.

**Figure 6.**
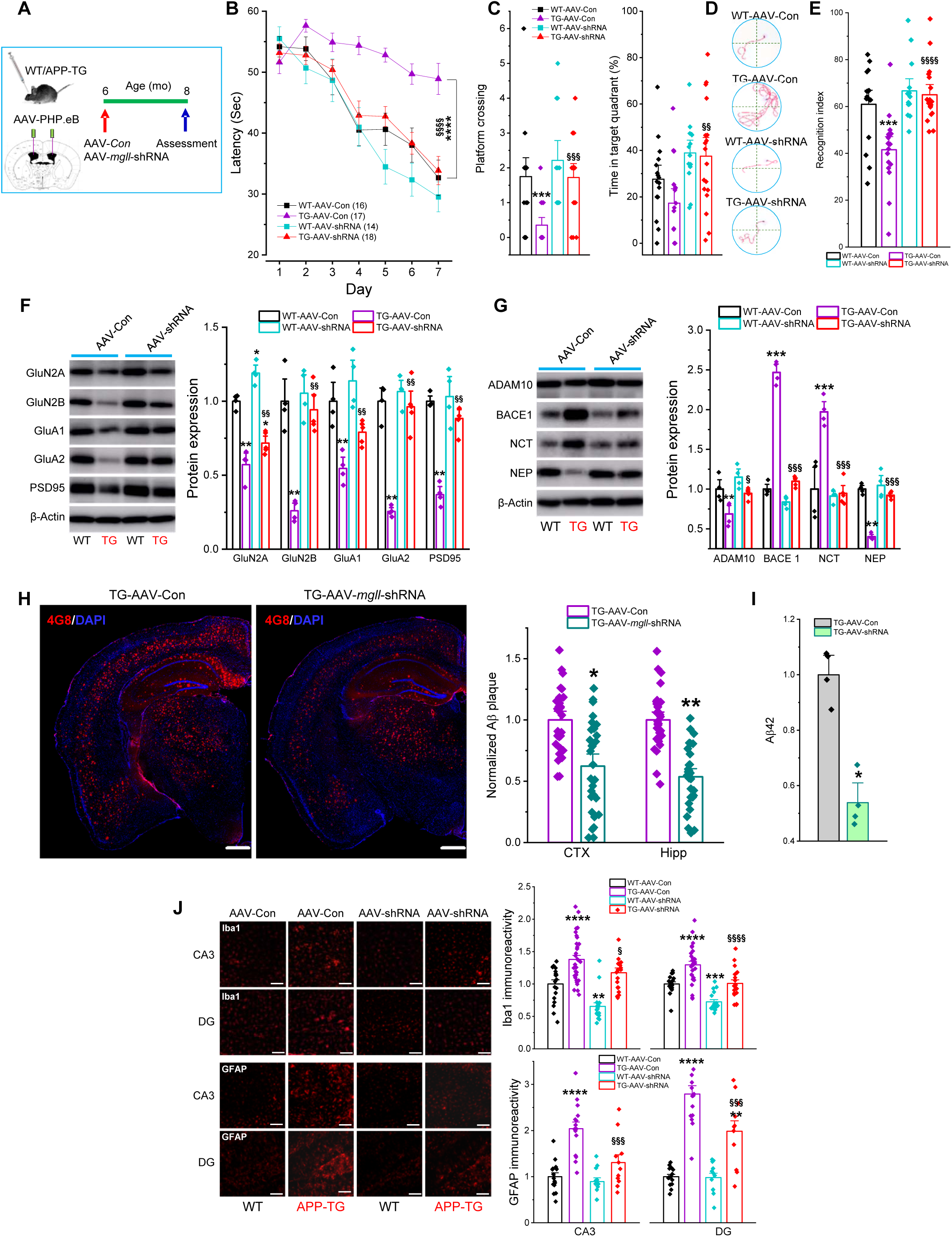
ICV delivery of AAV-*mgll*-shRNA vectors at a symptomatic stage reverses or delays AD progression in TG mice. (A) Schematic illustration of the protocol for ICV injection of AAV-*mgll*-shRNA vectors. (B) Learning acquisition in the MWM test. Data are presented as mean ± SEM. ****P<0.0001 vs WT-AAV-Con; ^§§§§^P<0.0001 vs TG-AAV-Con (Repeated measures ANOVA, n=14∼18). (C) Probe test. Data are presented as mean ± SEM (ANOVA with Bonferroni post-hoc test). (D) Representative swim paths during acquisition training on day 7. (E) NOR test. Data presented as mean ± SEM (ANOVA with Bonferroni post-hoc test, n=14∼18 animals/group). (F) Immunoblot analysis of hippocampal expression of glutamate receptor subunits and PSD95. Data are presented as mean ± SEM. **P<0.01 vs WT-AAV-Con, ^§§^P<0.01 vs TG-AAV-Con (Kruskal-Wallis test with Dunn’s post hoc test for multiple comparisons, n=4 animals/group). (G) Immunoblot analysis of hippocampal expression of Aβ-processing enzymes. Data are presented as mean ± SEM. *P<0.05, **P<0.01, ***P<0.001 vs WT-AAV-Con, ^§^P<0.05, ^§§§^P<0.001 vs TG-AAV-Con (Kruskal-Wallis test with Dunn’s post hoc test for multiple comparisons, n=4 animals/group). (H) Immunostaining of 4G8 immunoreactivity. Data are presented as mean ± SEM. *P<0.05, **P<0.01 vs TG-AAV-Con (Student’s t test, n=5 animals/group). Scale bars: 500 µm. (I) ELISA analysis of hippocampal Aβ42 levels. Data are presented as mean ± SEM. *P<0.05 (Mann-Whitney test, n=4 animals/group). (J) Immunostaining of GFAP and Iba1 reactivity in the brain. Data are presented as mean ± SEM. **P<0.01, ***P<0.001, ****P<0.0001 vs WT-AAV-Con, ^§§§^P<0.001, ^§§§§^P<0.0001 vs TG-AAV-Con (ANOVA with Bonferroni post-hoc test, n=5 animals/group). Scale bars: 100 µm.

## Discussion

In the present study, we provide compelling evidence that the beneficial effects of inhibiting 2-AG metabolism on neuropathology and on preserving synaptic and cognitive function in AD animals are cell type-specific: genetic deletion of MAGL in astrocytes, but not neurons, confers protection in TG mice. These protective effects are further corroborated by AAV-mediated silencing of MAGL in astrocytes, whereas overexpressing MAGL in astrocytes worsens neuropathology and accelerates synaptic and cognitive decline. Together with the elevated astrocytic MAGL expression observed in patients with AD and in TG mice in the present study, these findings suggest that astrocytic MAGL is a key pathogenic locus in the development of AD.

It is widely recognized that multiple factors contribute to the pathogenesis and progression of AD. This underscores the need to develop therapies that target molecules linked to several mechanisms or signaling pathways. MAGL is one such candidate, as it is the predominant enzyme hydrolyzing 2-AG in the brain ^14,15^, and 2-AG plays a critical role in maintaining brain homeostasis ^1,5–13,42^. On the other hand, some products of 2-AG hydrolysis are pro-inflammatory and neurotoxic prostaglandins (*e.g.*, PGE2) and leukotrienes (*e.g.*, LTB4) ^18–20^. Thus, a “*dual effect*” occurs when MAGL is inactivated, enhancing anti-inflammatory and neuroprotective 2-AG signaling, while simultaneously reducing proinflammatory and neurotoxic eicosanoids. Accumulated evidence suggests that neuroinflammation is a root cause of neurodegenerative diseases, including AD ^44,78,79^. Hence, resolving neuroinflammation is crucial for preventing development of AD or delaying progression of the disease ^1,32^.

Prior work from our lab and others has shown that systemic inactivation of MAGL, either through pharmacological inhibition or global deletion, alleviates Aβ pathology and improves synaptic and cognitive functions in AD model animals ^23,25,27,28^. Consequently, MAGL has been proposed as a therapeutic target for AD ^1,25,29,32^. However, accumulating evidence indicates that systemic MAGL inactivation may produce adverse effects in the brain and in peripheral tissues, including functional antagonism of the endocannabinoid system ^33,34^, impaired cerebellar fine motor coordination ^35^, increased incidence of lung adenocarcinoma ^36^, worsened cardiac function after acute myocardial infarction ^37^, compromised endothelial function and repair ^38^, and bone loss ^39^. These findings suggest that systemic MAGL inactivation may not be an ideal therapeutic strategy for AD. In particular, our recent study demonstrated that neuron-specific inactivation of MAGL leads to synaptic and cognitive impairments ^40^, indicating that intact neuronal 2-AG metabolism is essential for maintaining normal synaptic and cognitive function. In addition, overexpressing MAGL in astrocytes exacerbates neuroinflammation and neuropathology and accelerates synaptic and cognitive decline in TG mice, providing the first evidence that astrocytic 2-AG metabolism is directly linked to the development of AD, even though neurons produce the majority of 2-AG and astrocytes generate only a small fraction of total 2-AG in the brain ^17,42^. Moreover, proinflammatory prostaglandins are delivered predominantly from astrocyte-generated 2-AG hydrolyzed by MAGL ^17^. Thus, inactivating astrocytic MAGL would be expected to substantially reduce proinflammatory mediators ^17,42^.

The evidence presented in this study indicates that astrocyte-specific inactivation of MAGL represents a promising strategy for preventing or delaying the onset of AD, as well as for reversing disease progression, while avoiding synaptic and cognitive deficits caused by inhibiting 2-AG breakdown in neurons and the central and peripheral adverse effects associated with systemic MAGL inactivation, which leads to global blockade of 2-AG degradation.

Inactivation of astrocytic MAGL appears to underlie many of the beneficial effects observed in TG mice in this study. For example, synaptic abnormalities (including impaired synaptic plasticity, reduced expression of synaptic markers, and decreased synapse number) and cognitive deficits were prevented, and neuroinflammation, neurodegeneration, and Aβ pathology were alleviated in TG-aKO mice compared with TG mice, but not in TG-nKO mice. These protective effects of astrocytic MAGL inactivation in AD animals are likely mediated through pathways involving enhanced astrocytic 2-AG signaling and reduced eicosanoid signaling, leading to normalization of synapse-, immune/inflammation-, and AD-related DEGs; modulation of astrocyte-microglia, astrocyte-neuron, and microglia-neuron interactions to reduce neuroinflammation, neurodegeneration, and Aβ production while enhancing its clearance; preservation of synaptic structure, function, and plasticity.

Based on the data presented here, the protective effects of inhibiting 2-AG metabolism are clearly cell type-specific, and astrocyte-specific inactivation of MAGL emerges as an optimal therapeutic approach for AD. Because current pharmacotherapies cannot selectively target molecules in specific cell types, AAV-mediated gene replacement, gene silencing and gene editing offers a viable strategy with an excellent safety profile for CNS therapy ^80–82^. Indeed, both intrahippocampal and ICV delivery of AAV driven by a gfap promoter to silence astrocytic MAGL in TG mice at a presymptomatic stage prevented the onset of synaptic and cognitive decline, and mitigated neuroinflammation and Aβ pathology. Importantly, these protective effects were also observed when AAV-*mgll*-shRNA was delivered via ICV at a symptomatic stage. Together, these findings indicate that astrocyte-specific knockdown of MAGL can not only prevent AD development but also reverse or delay disease progression.

In the present study, we provide evidence that astrocyte-specific inactivation of MAGL represents a promising therapeutic strategy for the prevention and treatment of AD. Several approaches, including viral vectors, engineered nanoparticles, and extracellular vesicles (EVs), can serve as vehicles to deliver agents to defined cell populations. Here, we demonstrate that ICV delivery of AAV.PHP.eB vectors to silence astrocytic MAGL is a practical approach, as this vector exhibits high gene-transfer efficiency and is capable of crossing the blood-brain barrier (BBB). Although BBB penetration by AAV.PHP.eB in rodents is strain dependent, the vector has been shown to cross the BBB in non-human primates following intravenous administration ^76,77,83^. This suggests that, if translated into clinical settings, AAV.PHP.eB-based silencing of astrocytic MAGL could potentially be achieved through a single, minimally invasive intravenous injection as a therapeutic intervention for patients with AD.

## MATERIALS AND METHODS

### Animals

Cell type-specific *mgll* KO mice, including wild-type *mgll*^flox/flox^ (naïve), global knockout (tKO), astrocytic knockout (aKO), and neuronal knockout (nKO), were generated as described previously ^42^. These *mgll* KO lines were crossed with 5xFAD transgenic (TG) mice to generate 5xFAD TG cohorts with global (TG-tKO), astrocyte-specific (TG-aKO), or neuron-specific (TG-nKO) deletion of *mgll*, along with non-transgenic, non-knockout controls (WT). Because microglial MAGL contributes minimally to brain 2-AG hydrolysis, we did not generate 5xFAD mice lacking MAGL in microglia. The experiments were performed in a blinded fashion to prevent any scientific bias in the experiments. Animals were randomly assigned to experimental groups across genotypes. The number of animals per group was determined by power analysis, using 80% power, α = 0.05, and variance estimates based on our previously published data from similar experiments. Both male and female mice aged 8∼12 weeks were used in this study.

All animal procedures were performed in accordance with the U.S. Department of Health and Human Services Guide for the Care and Use of Laboratory Animals and were approved by the Institutional Animal Care and Use Committee of the University of Texas Health Science Center at San Antonio.

## METHOD DETAILS

### Western blots

Western blot analysis was performed to determine the expression of MAGL, PSD-95, ADAM10, BACE1, nicastrin (NCT), neprilysin (NEP), and glutamate receptor subunits (GluA1, GluA2, GluN2A, and GluN2B) in hippocampal tissues from WT and TG mice, as well as TG cohorts with cell type–specific mgll deletion. Hippocampal tissue was dissected, immediately homogenized in RIPA lysis buffer containing protease inhibitors, and incubated on ice for 30 min, followed by centrifugation at 10,000 rpm for 10 min at 4°C. Supernatants were fractionated on 4∼15% SDS-PAGE gels (Bio-Rad) and transferred onto PVDF membranes (Bio-Rad). Antibodies used to detect protein expression are listed in the Key Resources Table. Membranes were incubated with primary antibodies at 4°C overnight, washed, and then incubated with secondary antibody (goat anti-rabbit, 1:2,000; Cell Signaling) at room temperature for 1 hr. Proteins were visualized by enhanced chemiluminescence (ECL, Amersham Biosciences, UK). Band densities were quantified by densitometry using a GE/Amersham Imager 680 UV and normalized to total protein loading as determined by mouse anti–β-actin (Santa Cruz), as described previously ^27,42,84,85^.

### ELISA

ELISA analysis was performed as described previously ^27,85^. Whole hippocampi were lysed to generate total protein extracts. Tissues were homogenized in the lysis buffers provided with the respective ELISA kits for Aβ42 (AnaSpec, Inc.) and IL-1β (MilliporeSigma), and all procedures were performed according to the manufacturers’ instructions. Each kit included calibration (standard) curves, which were used to calculate absolute protein concentrations. Pilot experiments were conducted to optimize sample loading volumes and ensure that all measurements fell within the linear range of the standard curves. Absorbance was measured using a SpectraMax® iD3 plate reader (Molecular Devices). Technical replicates were included for each analyte to ensure accuracy and reproducibility.

### Hippocampal slice preparation

Hippocampal slices were prepared from mice as described previously ^25,27,42,84,85^. Briefly, after decapitation, brains were rapidly removed and placed in cold, oxygenated (95% O_2_, 5% CO_2_) artificial cerebrospinal fluid (ACSF) containing (in mM): 125.0 NaCl, 2.5 KCl, 1.0 MgCl_2_, 25.0 NaHCO_3_, 1.25 NaH_2_PO_4_, 2.0 CaCl_2_, 25.0 glucose, 3 pyruvic acid, and 1 ascorbic acid. Slices (350–400 µm thick) were cut and transferred to a holding chamber in an incubator containing ACSF at 36°C for 0.5∼1.0 h, then maintained in oxygenated ACSF at room temperature (22∼24°C) for >1.5 h before recording. Slices were subsequently transferred to a recording chamber and continuously perfused with 95% O_2_/5% CO_2_-saturated ACSF at ∼34°C.

### Electrophysiological recordings

Field EPSPs (fEPSPs) were recorded from hippocampal Schaffer collateral-CA1 synapses in response to 0.05 Hz stimulation using an Axoclamp-2B and Axoclamp 900A amplifiers, as described previously. Recording pipettes were pulled from borosilicate glass using a micropipette puller (Sutter Instruments) and filled with ACSF (resistance ∼4 MΩ). As previously described ^42,85^, long-term potentiation (LTP) at CA3-CA1 synapses was induced by high-frequency stimulation (HFS) consisting of three trains of 100 Hz stimulation (1-second duration each) delivered with a 20-second inter-train interval. EPSP potentiation following HFS was normalized to the pre-stimulation baseline.

### Golgi–Cox staining

Golgi-Cox staining was used to visualize dendritic spines of hippocampal neurons as previously described ^40,86^, with minor modifications. WT and TG mgll-deletion cohorts were transcardially perfused with ice-cold saline for 5 minutes under anesthesia. Brains were dissected and processed using the Golgi-Cox Impregnation & Staining System according to the manufacturer’s instructions (superGolgi Kit, Bioenno Tech, LLC; Cat# 003010). After impregnation, coronal sections (100–200 µm) were cut on a vibratome, mounted on gelatin-coated slides, and stained. Images were acquired using a Zeiss Imager II deconvolution microscope with SlideBook 6.0 software. For dendritic spine quantification, z-stack images were collected at 0.1-µm steps to generate sequential optical sections for spine counting and morphological analysis on 3D reconstructions using a 100× oil-immersion objective. NeuronStudio (Version 0.9.92; http://research.mssm.edu/cnic/tools-ns.html, CNIC, Mount Sinai School of Medicine) was used to reconstruct and analyze dendritic spines, as described previously.

### Transmission electron microscopy (TEM)

For TEM experiments, all tissues were processed using freshly prepared solutions on the day of perfusion, as described previously ^40^. Briefly, animals were anesthetized and transcardially perfused with normal saline, followed by 2.5% glutaraldehyde/4% paraformaldehyde EM fixative (in 0.16 M NaH_2_PO_4_/0.11M NaOH buffer, pH 7.2∼7.4) for 30 min. After perfusion, whole carcasses were post-fixed for at least 1 week in the same EM fixative. The hippocampus was then dissected and incubated overnight in 0.1 M sodium cacodylate buffer, followed by incubation in a 2% OsO_4_ solution and gradient ethanol dehydration. Samples were incubated in propylene oxide, left in 100% PolyBed resin for 36 hours, and embedded in flat molds at 55°C for 36 hours. After embedding, the molds were processed, sectioned at a thickness of 90 nm, and imaged on a JEOL 1400 electron microscope in the Electron Microscopy Lab at UT Health San Antonio. Synapses were identified by the presence of presynaptic vesicles, a clearly defined synaptic cleft, and postsynaptic densities. The number of synapses was manually counted and quantified in each image, as described previously.

### AAV injection

The AAV vectors used in the present study were obtained from Vector Biolabs: AAV5-gfap-eGFP-*mgll*-shRNAmir(2) (1.7 x 10^13^ GC/ml), AAV5-gfap-h*mgll*-eGFP (6.0 x 10^12^ GC/ml), AAV5-gfap-eGFP (2.3 x 10^13^ GC/ml), AAV/PHP.eB-gfap-eGFP-mgll-shRNAmir(2) (1.8 x 10^13^ GC/ml), and AAV/PHP.eB-gfap-eGFP (1.2 x 10^13^ GC/ml). WT or 5xFAD mice were anesthetized with ketamine/xylazine (200/10 mg/kg) and placed in a stereotaxic frame. AAV5-gfap-eGFP-mgll-shRNAmir(2), AAV5-gfap-hmgll-eGFP, or AAV5-gfap-eGFP was stereotaxically injected into both hippocampi at the following coordinates: AP, -2.3; ML, ±2.1; DV, -2.0, as described previously ^27,42,85,86^. AAV/PHP.eB-gfap-eGFP-mgll-shRNAmir(2) and AAV/PHP.eB-gfap-eGFP were stereotaxically injected into the right and left lateral ventricles at the following coordinates: AP, -0.5; ML, ±1.2; DV, -2.1 from the brain surface. After AAV delivery, the incision was closed with sterile silk sutures. Mice were kept in a heated cage for at least 1 hour and then returned to their home cages.

### Single-cell/nucleus sample preparation

Single-cell suspensions from WT, TG, TG-tKO, TG-aKO, and TG-nKO mice were prepared using an Adult Brain Dissociation Kit (MACS Miltenyi Biotec, Cat# 130-107-677) according to the manufacturer’s instructions, with some modifications, as previously described ^40,42,47^. Single-nucleus suspensions from the animals were prepared using a protocol previously described with some modification. Briefly, frozen hippocampi from five animals for each genotype were homogenized in ice-cold lysis buffer on ice. The resulting suspension was filtered through a 20 μm filter to remove debris and centrifuged at 500 g for 5 minutes at 4°C. Nuclei were washed and filtered twice with nuclei wash buffer. Finally, the pellets were carefully resuspended in a suitable volume of nuclei buffer to achieve a concentration of 500-1,000 nuclei/μl for subsequent capture, as previously described.

### Single-cell/nucleus RNA sequencing library preparation

Single-cell or nucleus suspensions were loaded into the 10x Genomics Chromium microfluidic chips with the intention of capturing 8,000 to 10,000 cells within individual Gel Beads-in-emulsion (GEM). Within the GEMs, cell lysis occurred, and RNA was reverse transcribed using poly(dT) priming, during which Cell Barcodes and Unique Molecular Identifiers (UMIs) were incorporated into the cDNA. The prepared libraries, following the 10x Genomics 3’ Gene Expression v3 protocol, were sequenced using the Illumina NovaSeq 6000 system at the Genome Sequencing Facility (GSF) of Greehey Children’s Cancer Research Institute at UT Health San Antonio.

### Single nucleus RNA-Seq data analysis

Data analysis involved Cell Ranger (v4.0) and Seurat (v5.0.3) in R (v4.2.1). CellBender (version 0.2.0) was used to remove ambient RNA before downstream analysis ^87^. Low-quality cells/nuclei were excluded using thresholds (>30% mitochondrial or >20% ribosomal gene expression). Genes detected in <10 nuclei were removed, with per-nucleus thresholds set at 400–6,000 genes and ≤35,000 UMIs. Data normalization (scale factor 10,000) and identification of variable features (5,000 genes via ‘vst’) preceded PCA-based dimensionality reduction (top 25 PCs). Clustering (resolution 2) and UMAP visualization were performed.

After checking the quality of the snRNA-seq data, we used 24,614 cells for further analysis, with an average of 3613 genes. We worked with R packages like Seurat to group and label the cells using well-known brain marker genes (Supplementary Figure 1A) Cell-type annotation of mouse hippocampal cells relied on known markers in: *syt1*/*gad1*/*gad2* (inhibitory neurons), *syt1*/*slc17a7*/*cpne4*/*prkcb* (CA3 pyramidal neurons), *syt1*/*slc17a7*/*arhgap12*/*fibcd1* (CA1 pyramidal neurons), *syt1*/*slc17a7*/*stxbp6*/*prox1* (DG granule cells), *gjb6*/*gja1* (astrocytes), *c1qa*/*cx4cr1* (microglia), and *plp1*/*mog* (oligodendroglia). Cell-type annotation of human hippocampal cells relied on known markers (Supplementary Figure 2E): *SYT1*/*GAD*1/*GAD2* (inhibitory neurons), *SYT1*/*CAMK2A*/*CPNE4* (CA3 pyramidal neurons), *SYT1*/*CAMK2A*/*PEX5L* (CA1 pyramidal neurons), *SYT1*/*CAMK2A*/*STXBP6* (DG granule cells), *AQP4*/*SLC1A2* (astrocytes), *CD74*/*CSF1R* (microglia), and *PLP1*/*MBP* (oligodendroglia). In general, differentially expressed genes (DEGs) were calculated by comparing TG mice to WT mice, and the crossbred mice (TG-tKO, TG-aKO, TG-nKO) to TG mice. DEGs were identified using Seurat’s FindMarkers (Wilcoxon test; P < 0.05, |log2FC| > 0.1), with synaptic and neurogenesis-related genes filtered using SYNGO and MANGO databases. Volcano plots were produced by OriginLab 2024. Heatmaps were plotted with normalized expression values produced by scRNAtoolVis package.

### Gene Ontology analysis

GO analyses were generated via ClusterProfiler (version 4.10.1) and Org.Mm.eg.db (version 3.18.0) with default parameters.

### Gene network analysis

Significantly expressed synaptic or immune/inflammation-related DEGs from different cell types in the TG-aKO group (compared with the TG group) were uploaded to the STRING database to construct gene interaction networks. The primary interaction source was set to “Experiments.” If no experimentally verified interactions were available between genes, other sources (excluding “Textmining”) were used to generate the networks. In this study, synaptic gene networks were constructed using the “Experiments” source, while immune/inflammation-related gene networks were constructed using the “Experiments,” “Database,” “Co-expression,” “Neighborhood,” “Gene Fusion,” and “Co-occurrence” sources. The resulting networks were exported as TSV files and manually refined in Cytoscape (version 3.10.4).

### Novel object recognition test

The novel object recognition (NOR) test was performed to assess memory retention as described previously ^26,40,86^. Briefly, the NOR test was conducted in a 30 × 30 × 30 cm square open field. The animals were first allowed to acclimate to the testing environment (habituation). The NOR assay then comprised two 10-minute stages separated by a 4-hour intertrial interval. In the training stage, animals were presented with two identical objects. In the test stage, animals were presented with one familiar object (from training) and one novel object. Exploration was defined as the time spent with the head oriented toward and within 2 cm of an object. Object exploration during the test stage was quantified using the EthoVision video-tracking system (Noldus). The recognition index (RI) was calculated based on the following equation: RI =T_N_/T_N_+T_F_), where T_N_ is the exploration time devoted to the novel object and T_F_ is the exploration time for the familiar object, as described previously ^26,40^.

### Morris water Maze test

The classic Morris water maze (MWM) test was used to assess spatial learning and memory, as described previously ^27,42,85,86^. A circular water tank (120 cm in diameter and 75 cm in height) was filled with water rendered opaque using non-toxic white paint. A round platform (15 cm in diameter) was submerged 1 cm below the water surface at the center of one quadrant of the tank.

Mice received acquisition training for 7 consecutive days, with 4 trials per day. For each trial, the mouse was released from the wall of the tank and allowed up to 60 seconds to locate the hidden platform, on which it remained for 5 seconds once found. For each training session, the starting position and the sequence of the four quadrants from which the mouse was released were randomly varied across sessions and across animals.

Mouse movement in the pool was recorded by a video camera, and task performance, including swim path, swim speed, and time spent in each quadrant, was analyzed using the EthoVision video tracking system (Noldus, version 17). A probe trial was conducted 24 hours after completion of acquisition training, during which the platform was removed, and performance was recorded for 60 seconds.

### Data and code availability

All materials are readily available from the corresponding author on reasonable request. This paper does not report original code. All other materials can be obtained from commercial vendors. Sing-nucleus RNA sequencing data have been deposited at the NCBI Gene Expression Omnibus (GEO) under accession number GSE311467. All other data reported in this paper will be shared by the lead contact upon request.

### Quantification and statistical analysis

Data are presented as mean ± S.E.M. Unless stated otherwise, parametric tests (*e.g.,* Student’s t-test, one- or two-way analysis of variance followed by appropriate post hoc tests) were used for statistical comparisons. Non-parametric alternatives (*e.g.,* Mann-Whitney for two groups and Kruskal-Wallis followed by Dunn’s tests for multiple comparisons) were used when normality assumptions were not met. Differences were considered statistically significant at P < 0.05.

## Supporting information

Related to Figure-1

Related to Figure-3

Related to Figure-3

Related to Figure-4

Related to Figure-5

## ACKNOWLEDGMENTS

This work was supported by National Institutes of Health grants RF1NS076815 and RF1AG081362 (to C.C.) and by startup funds from UT Health San Antonio, Joe R. & Teresa Lozano Long School of Medicine (to C.C.). RNA sequencing data was generated in the Genome Sequencing Facility at UT Health San Antonio, which is supported by UT Health San Antonio, NIH Shared Instrument grant S10OD030311, and CPRIT Core Facility Award (RP220662). The authors thank NIH/Harvard Brain Tissue Resource Center for providing human hippocampal tissues, Ms. Barbara Hunter for her technical support with TEM, and Mr. Jack Hashem and Ms. Anastassia R. Nelson for their assistance with animal care.

## AUTHOR CONTRIBUTIONS

C.C. and J.Z. conceived the project and designed the experiments; L.S., M.H., J.L., D.Z., F.G., M.P., J.Z., and C.C. performed the experiments and/or analyzed the data; C.C. and J.Z. supervised the work and C.C. wrote the manuscript.

