## Supplementary material for "Selective Inactivation of Astrocytic Monoacylglycerol Lipase for Alzheimer’s Disease Therapy": Related to Figure-1

### Slide 1
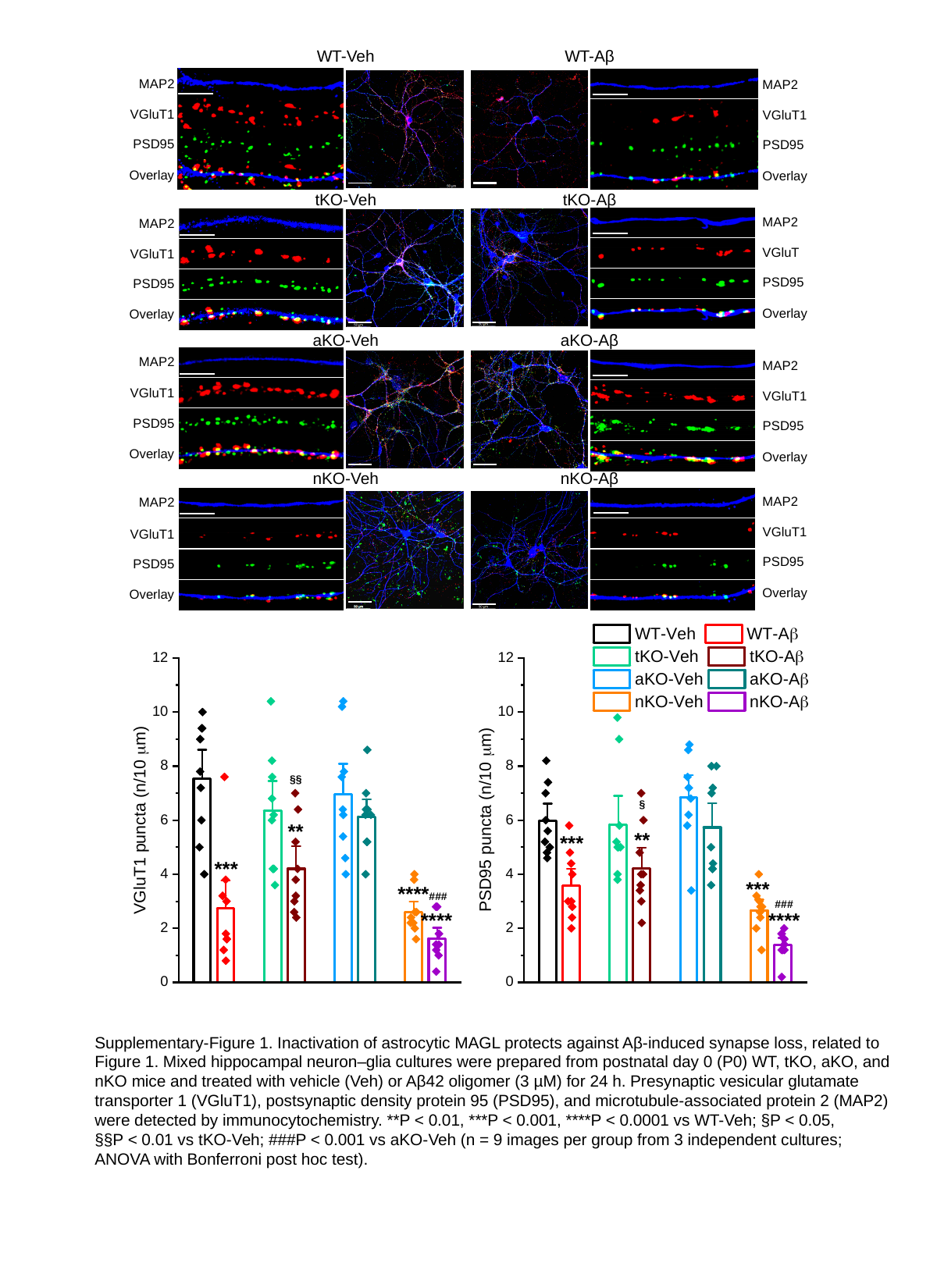

WT-Veh
WT-Aβ
MAP2
MAP2
VGluT1
VGluT1
PSD95
PSD95
Overlay
Overlay
tKO-Veh
tKO-Aβ
MAP2
MAP2
VGluT
VGluT1
PSD95
PSD95
Overlay
Overlay
aKO-Veh
aKO-Aβ
MAP2
MAP2
VGluT1
VGluT1
PSD95
PSD95
Overlay
Overlay
nKO-Veh
nKO-Aβ
MAP2
MAP2
VGluT1
VGluT1
PSD95
PSD95
Overlay
Overlay
5 um
50 um
Aβ (3 uM)
Supplementary-Figure 1. Inactivation of astrocytic MAGL protects against Aβ-induced synapse loss, related to
Figure 1. Mixed hippocampal neuron–glia cultures were prepared from postnatal day 0 (P0) WT, tKO, aKO, and
nKO mice and treated with vehicle (Veh) or Aβ42 oligomer (3 µM) for 24 h. Presynaptic vesicular glutamate
transporter 1 (VGluT1), postsynaptic density protein 95 (PSD95), and microtubule-associated protein 2 (MAP2)
were detected by immunocytochemistry. **P < 0.01, ***P < 0.001, ****P < 0.0001 vs WT-Veh; §P < 0.05,
§§P < 0.01 vs tKO-Veh; ###P < 0.001 vs aKO-Veh (n = 9 images per group from 3 independent cultures;
ANOVA with Bonferroni post hoc test).
