## Supplementary material for "Selective Inactivation of Astrocytic Monoacylglycerol Lipase for Alzheimer’s Disease Therapy": Related to Figure-3

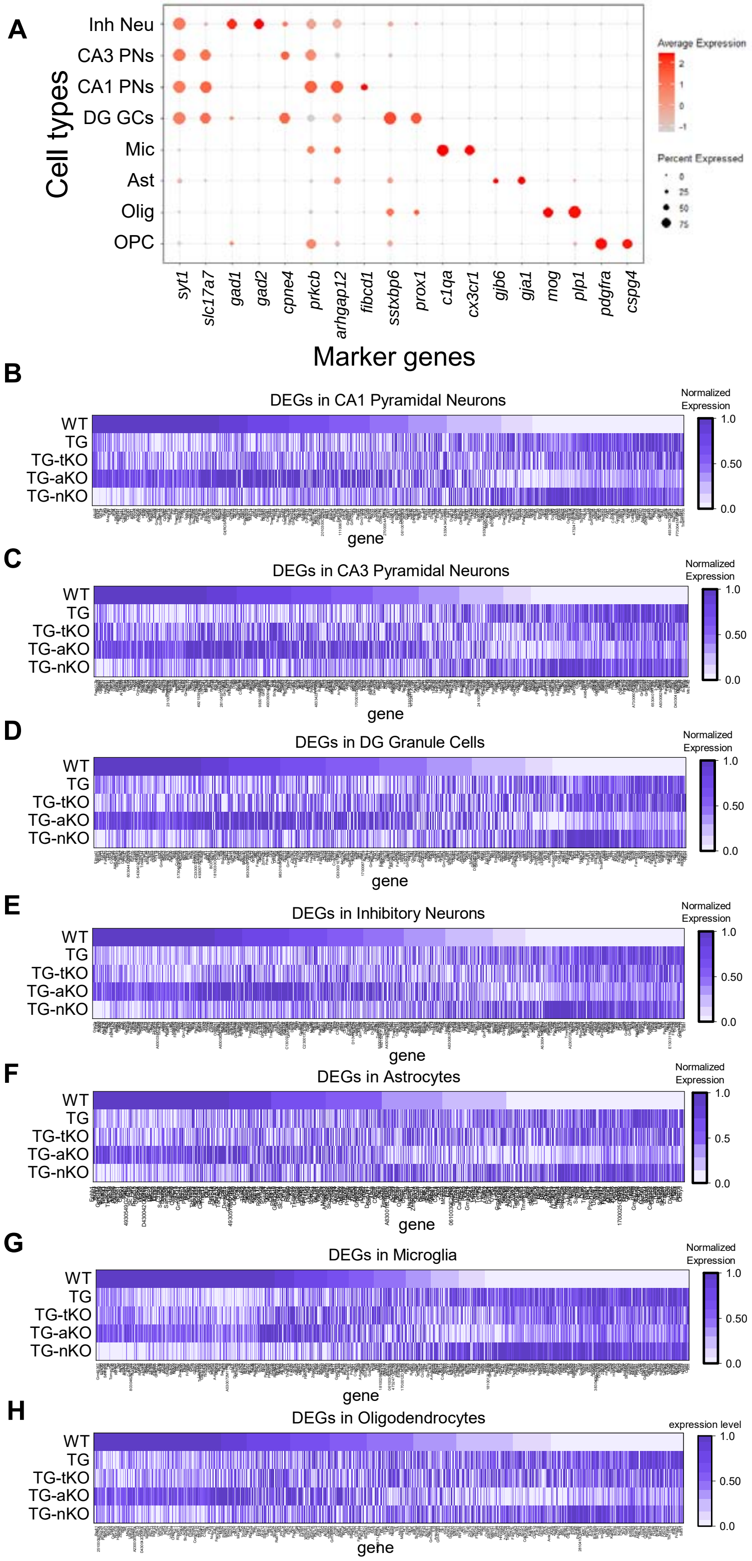


Supplementary Figure 2. Summary of cell type-specific differentially expressed gene (DEG) expression levels, related to Figure 3. Dot plot showing the expression of cell type-specific markers used to identify major hippocampal cell populations in mice. (B~H) Heatmaps showing normalized expression levels of DEGs in CA1 pyramidal neurons (PNs) (B), CA3 PNs (C), dentate gyrus granule cells (DG GCs) (D), inhibitory neurons (Inh Neu) (E), astrocytes (Ast) (F), microglia (Mic) (G), and oligodendrocytes (Olig) (H) in TG mice compared with wild-type controls.
