## Supplementary material for "Selective Inactivation of Astrocytic Monoacylglycerol Lipase for Alzheimer’s Disease Therapy": Related to Figure-3

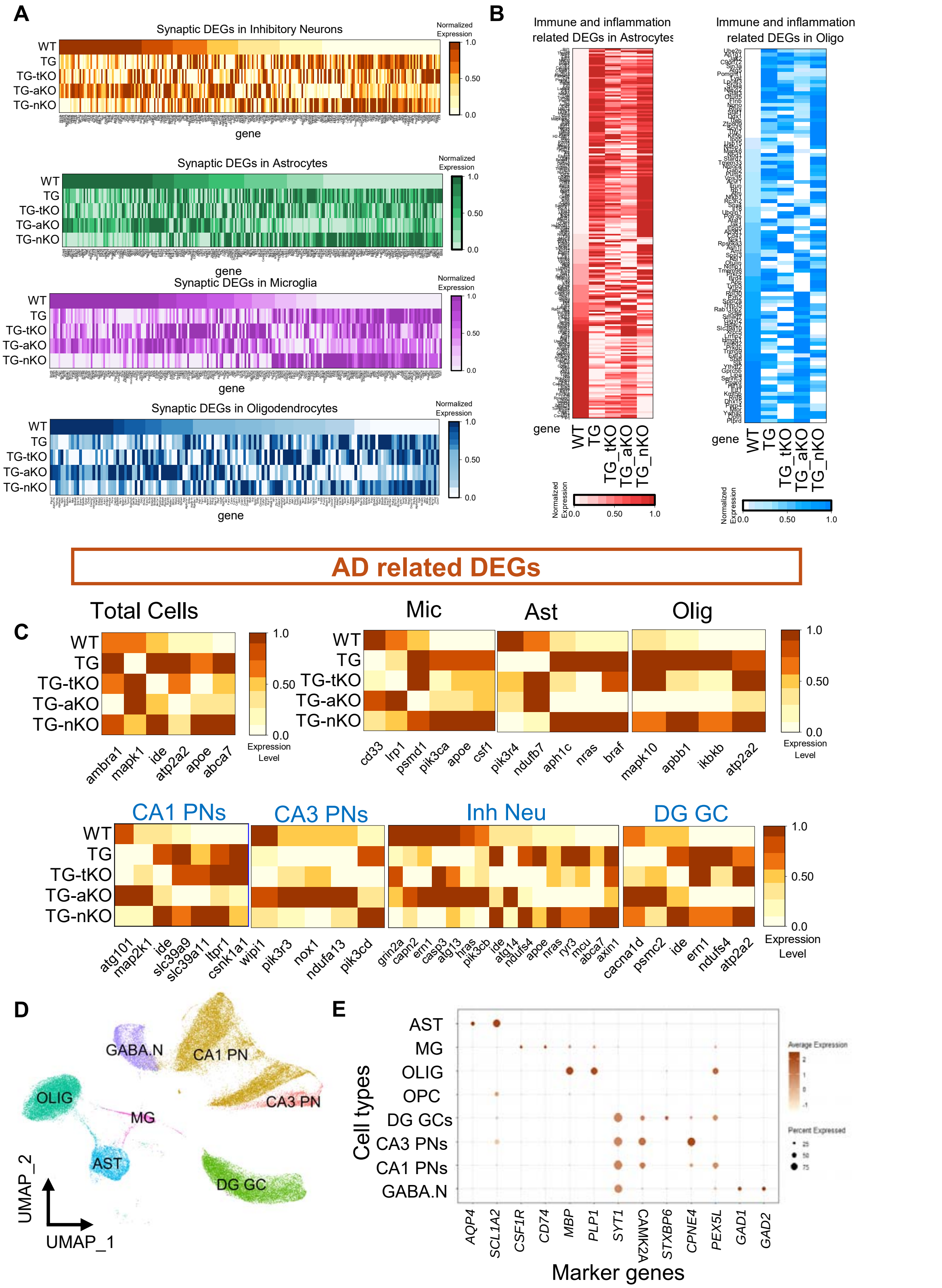


Supplementary Figure 3. Heatmaps showing normalized expression levels of synaptic, immune/inflammation-related, and AD-related genes across different cell types, related to Figure 3. (A) Heatmaps illustrating expression patterns of synaptic genes in inhibitory neurons (Inh Neu), astrocytes (Ast), microglia (Mic), and oligodendrocytes (Olig) across different transgenic groups. (B) Heatmaps showing normalized expression levels of immune/inflammation-related DEGs in Olig. (C) Heatmaps displaying normalized expression levels of AD-related DEGs in total cells, Ast, Mic, Olig, CA1 pyramidal neurons (PNs), CA3 PNs, dentate gyrus granule cells (DG GCs), and Inh Neu. Dark colors indicate high expression levels, whereas light colors indicate low expression levels. (D) t-SNE plot of snRNA-seq datasets re-analyzed from hippocampi of AD patients and normal controls. (E) Dot plot showing the expression of cell type-specific markers used to identify major hippocampal cell populations in AD patients and normal controls.
