## Supplementary material for "Selective Inactivation of Astrocytic Monoacylglycerol Lipase for Alzheimer’s Disease Therapy": Related to Figure-4

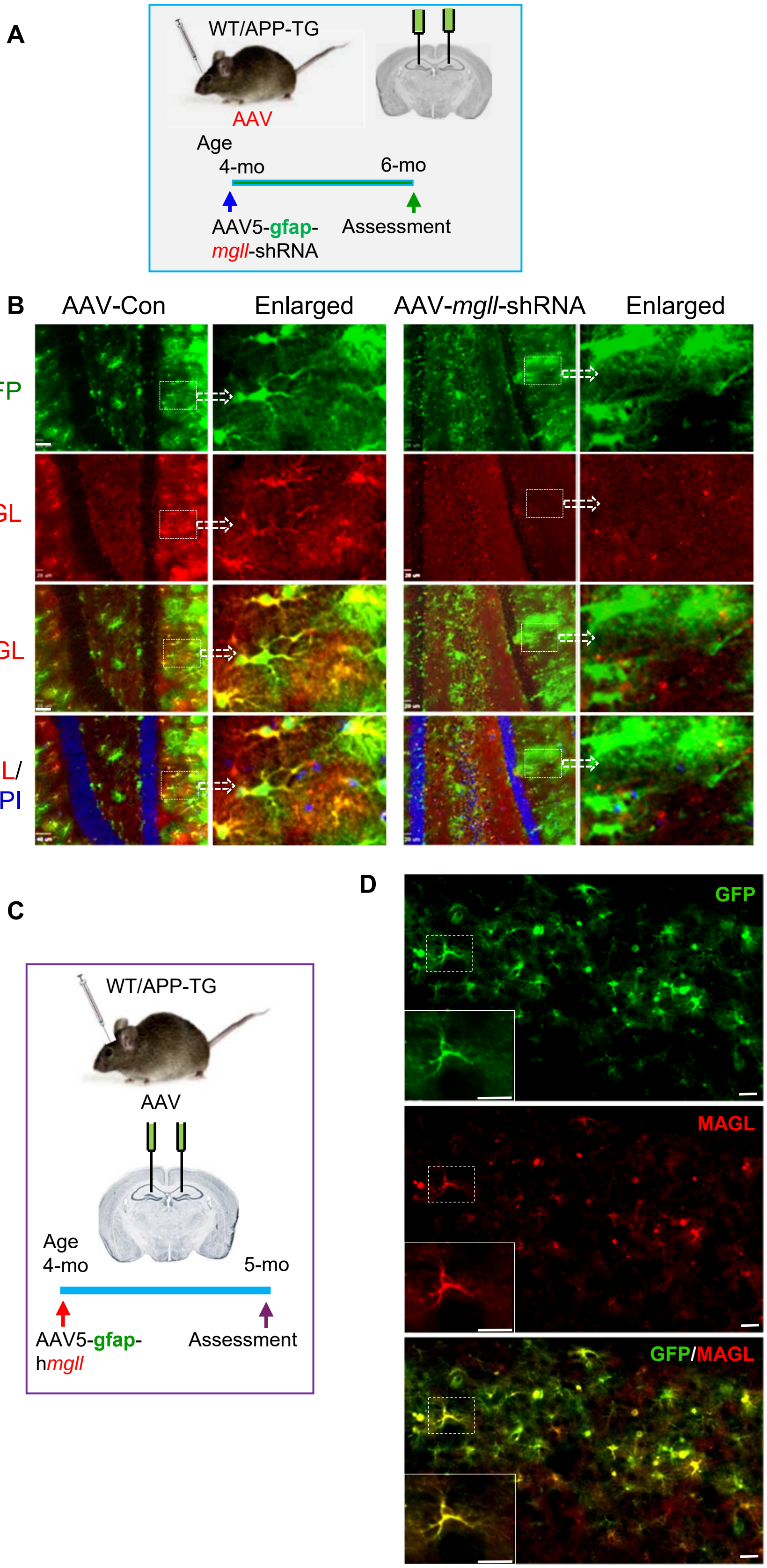


Supplementary Figure 4. Intrahippocampal injection of AAV5-*mgll*-shRNA or AAV5-h*mgll*, related to Figure-4 (**A**) Schematic illustration of the protocol for intrahippocampal AAV-*mgll*-shRNA injection. (**B**) Immunostaining showing MAGL expression in the hippocampus following AAV-*mgll*-shRNA injection. (**C**) Schematic illustration of the protocol for intrahippocampal AAV-h*mgll* overexpression. (**D**) Immunostaining showing MAGL overexpression in the hippocampus following AAV-h*mgll* vector injection.
