## Supplementary material for "Selective Inactivation of Astrocytic Monoacylglycerol Lipase for Alzheimer’s Disease Therapy": Related to Figure-5

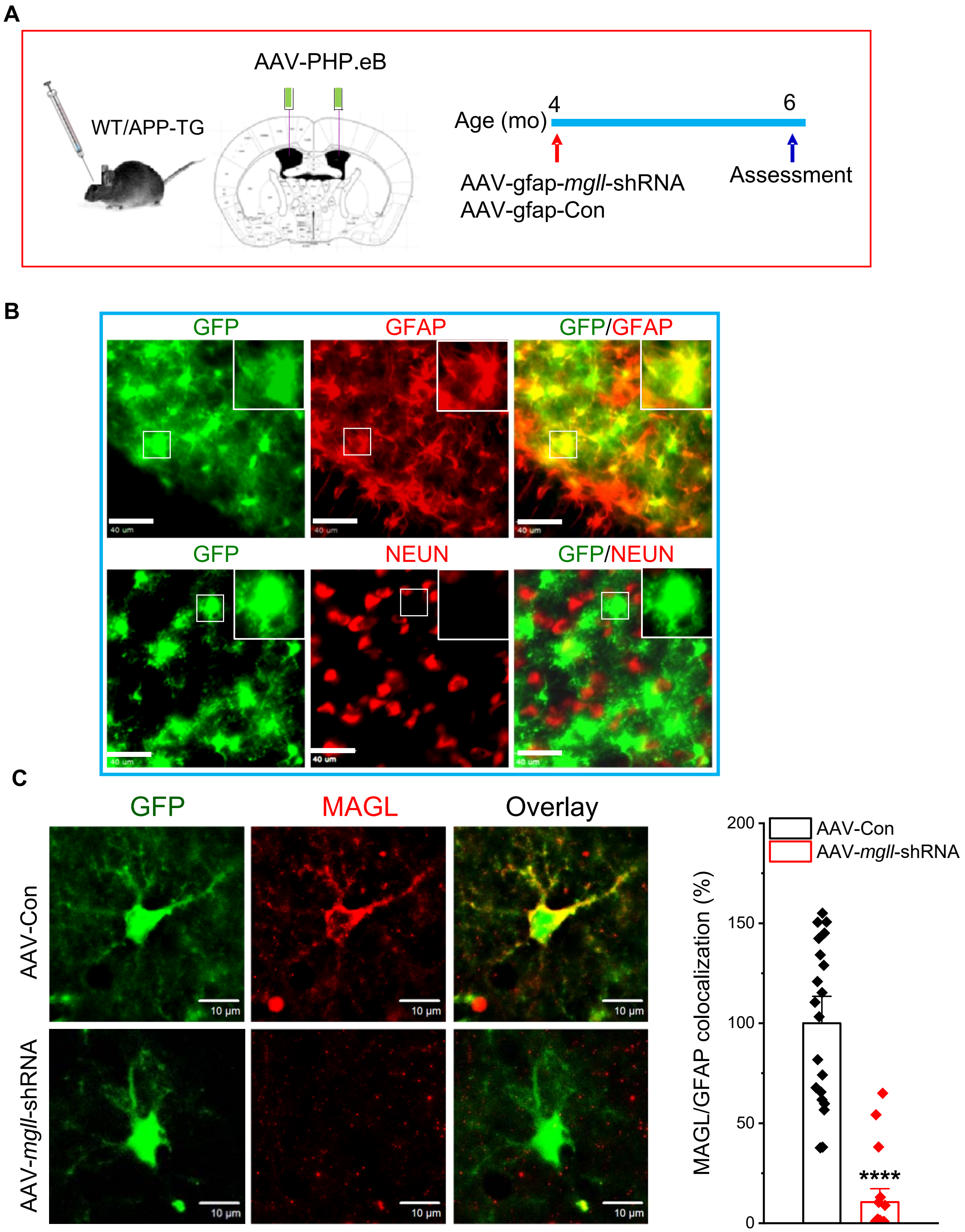


Supplementary Figure 5. Intracerebroventricular (ICV) injection of AAV.PHP.eB-gfap-GFP-*mgll*-shRNA, related to Figure 5. (**A**) Schematic illustration of the protocol for ICV injection of AAV-*mgll*-shRNA. (**B**) Immunostaining showing colocalization of GFP with GFAP, but not with NeuN following AAV vector injection. (**C**) Immunostaining showing reduced MAGL expression in astrocytes expressing the AAV-*mgll*-shRNA vector. Data represented as mean ± SEM ****P<0.0001 (ANOVA with Bonferroni post-hoc test, n=4 animals/group).
